# Functional stability and recurrent STDP in rhythmogenesis

**DOI:** 10.1101/2025.03.04.641421

**Authors:** Gabi Socolovsky, Maoz Shamir

**Affiliations:** Department of Physics, Faculty of Natural Sciences, Ben-Gurion University of the Negev, Be’er-Sheva, Israel; The School of Brain Sciences and Cognition, Ben-Gurion University of the Negev, Be’er-Sheva, Israel; Department of Physiology and Cell Biology, Faculty of Health Sciences, Ben-Gurion University of the Negev, Be’er-Sheva, Israel

## Abstract

Synapses in the nervous system show considerable volatility, raising the question of how the brain maintains functional stability despite continuous synaptic motility. Previous studies have suggested that functionality may be maintained by an ongoing process of activity-dependent plasticity. Here, we address this question in the context of rhythmogenesis. Specifically, we investigated the hypothesis that rhythmic activity in the brain can develop and be stabilized via activity-dependent plasticity in the form of spike-timing-dependent-plasticity (STDP), thus extending our previous work that demonstrated rhythmogenesis in a toy model of two effective neurons. We examined STDP dynamics in large recurrent networks in two stages. We first derived the effective dynamics of the order parameters of the synaptic connectivity. Then, a perturbative approach was applied to investigate stability. We show that for a wide range of parameters, STDP can induce rhythmogenesis. Moreover, STDP can suppress synaptic fluctuations that disrupt functionality. Interestingly, STDP can channel fluctuations in the synaptic weights into a manifold in which the network activity is not affected, thus maintaining functionality while allowing a subspace in which synaptic weights can be widely distributed.

## I. INTRODUCTION

It is widely accepted that our sensory perception as well as planned motor commands are represented in the brain by the dynamical state of neural activity. Synaptic connections shape the space of this activity and hence the functionality of the brain. However, empirical findings report that the strength of synaptic connections; i.e., synaptic weights, is highly volatile [1–8]. Synaptic weights have been shown to fluctuate by about 50 percent over the course of several days [6, 7]. Moreover, activity-independent fluctuations in the synaptic weights with a similar order of magnitude have also been observed [4, 5, 7, 8]. This raises the question of how functionality can be maintained in the face of constant synaptic motility.

Here, we address the question of functional stability within the context of rhythmogenesis. Rhythmic activity has been widely observed in the central nervous systems of a diverse range of animals and has been linked to various cognitive tasks [9–17]. Numerous computational studies have investigated models that can generate this type of rhythmic activity [9, 18–26]. In these models, for rhythmic activity to emerge, the synaptic weights must be within a certain range of values. Thus, in the context of rhythmogenesis, the question of functionality relates to into the mechanism that ensures that the synaptic weights develop to the target range and stabilizes them. Activity-dependent synaptic plasticity, and in particular Spike-Timing-Dependent Plasticity (STDP), has been suggested as a potential mechanism to maintain functionality [27–31]. In STDP, the synaptic weight is modified according to the time difference between the pre- and post-synaptic firings [32–36]. Various STDP learning rules have been observed across multiple brain regions and in numerous animal species [33, 34, 37–42].

We previously showed, as a proof of concept, that STDP can lead the development of synaptic weights into the required range for rhythmic activity [43]. However, this was done in the framework of a minimal model, which is equivalent to a network having one excitatory and one inhibitory neuron with only reciprocal connections. Thus, it remains unclear whether STDP can provide a mechanism for rhythmogenesis in large networks with other types of connectivity and motility of individual synapses.

To address these questions, we examined rhythmogenesis within the framework of a rate model introduced by Roxin et al. [23] (see also [24, 25, 44]), which comprises large excitatory and inhibitory populations that incorporates various types of synaptic connections. Specifically we focused on the STDP dynamics of the interconnections, since these connections are the essential synapses for generating the rhythmic activity (fig. 1).

**FIG. 1:**
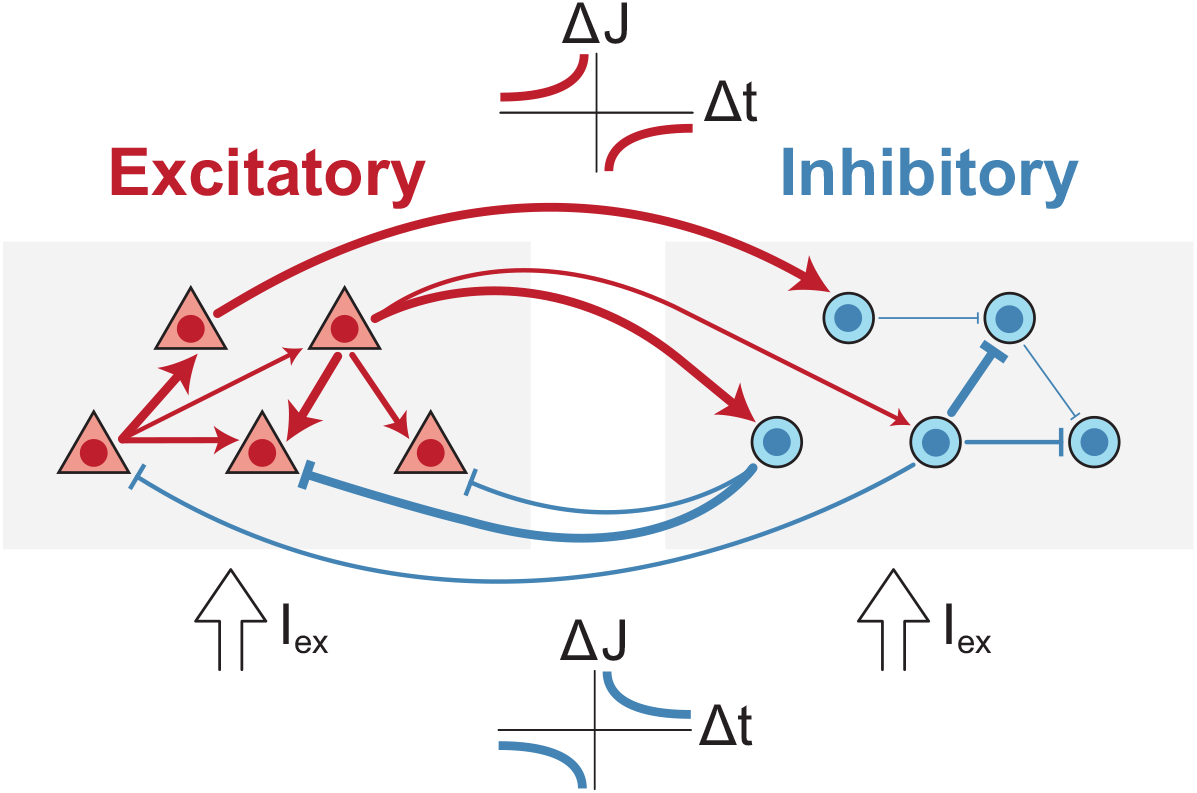
Network architecture. The network consists of an excitatory population (red) and an inhibitory population (blue). Neurons are fully connected both within and between populations, with synaptic strengths represented by varying synapse thicknesses. A Hebbian STDP learning rule is applied to inhibitory-to-excitatory (I-to-E) connections, and an anti-Hebbian rule is applied to excitatory-to-inhibitory (E-to-I) connections. All neurons receive the same constant external input.

This paper is structured as follows. In section II we define the dynamical model of the neural responses. In II A, we derive the phase diagram of the network in the case where all synapses of the same type are equal, and identify the mean (across the population) synaptic weights as order parameters of the system. Then, in II B, we extend the analysis by incorporating small fluctuations in the synaptic weights around the order parameters, and compute the cross-correlations in the neural responses. In Section III we introduce the STDP dynamics. In III A we define the STDP learning rules used in this study and develop the dynamical equations for synaptic weights in the limit of slow learning. In our approach, STDP dynamics is studied in two stages. First, we study the induced STDP dynamics of the order parameters, III B. Then, the STDP dynamics of fluctuations around the order parameters are analyzed using a perturbative approach, III C. Finally, we summarize our results and discuss our work in the context of maintaining functional stability in the face of constant synaptic motility, and theoretical approaches for studying recurrent STDP.

## II. FIRING RATE DYNAMICS

We model a system of *N*_*E*_ excitatory (*E*) neurons and *N*_*I*_ inhibitory (*I*) neurons (fig. 1). The spiking activity of each neuron follows inhomogeneous Poisson statistics, where 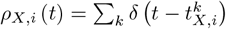 is the spike train of neuron *i* in population *X* ∈ {*E, I*}, expressed as the sum of Dirac’s delta functions at times 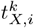. Following Roxin et al. [23], the instantaneous firing rate of neuron *i* in population *X, m*_*X,i*_ (*t*) = ⟨*ρ*_*X,i*_ (*t*)⟩, where ⟨…⟩ denotes ensemble averaging, is governed by:

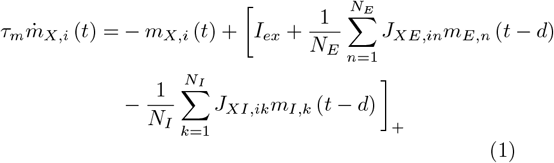

where *J*_*XY,ij*_ is the synaptic connection from neuron *j* in population *Y* ∈ {*E, I*} to neuron *i* in population *X*. The characteristic time of the neuronal response is *τ*_*m*_, and *I*_*ex*_ is an external input. The parameter *d* is the delay in the neural response to its input (see [26] for a discussion of various physiological sources for the delay). The function [*x*]_+_ denotes the threshold linear function; i.e., [*x*]_+_ = *H* (*x*) *x*, where *H*(*x*) is the Heaviside function. Unless stated otherwise, we measure time in units of the characteristic time of the neural response; i.e., *τ*_*m*_ = 1, and the neural responses are measured relative to the external input, i.e., *I*_*ex*_ = 1.

Note that although this model contains autapses, *J*_*XX,ii*_, their contribution to the total synaptic input to each neuron is negligible for large networks, *N*_*E*_, *N*_*I*_ ≫1.

In the following subsections we analyze the neural dynamics in the case of uniform synapses and extend these results to small fluctuations around the uniform synapses solution.

### A. Homogeneous network and order parameters

In the case of uniform synaptic coefficients, *J*_*XY,ij*_ = *J*_*XY*_ ∀ *i, j*, a uniform solution, in which *m*_*X,i*_(*t*) = *m*_*X*_ (*t*) ∀*i*, exists and is stable, see also [23, 26, 43]. In this case eq. (1), is reduced to

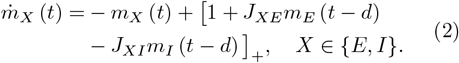

The range of the synaptic weights we chose to investigate revealed four global states of activity (see detailed derivation in Appendix A 1). Specifically, since we were interested in rhythmic activity that results from E-I interactions, we assumed moderate intra-connection strengths; i.e., *J*_*EE*_, *J*_*II*_ *<* 1.

Depending on the values of the synaptic weights, eq. (2) exhibits different types of behaviors, figs. 2a and 2b. For sufficiently strong I-to-E inhibition, *J*_*EI*_ *>* 1 + *J*_*II*_, the activity of the excitatory population can be fully suppressed by inhibition and the system converges to a fixed-point in which Only Inhibition (OI) is active,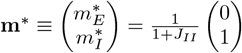.

**FIG. 2:**
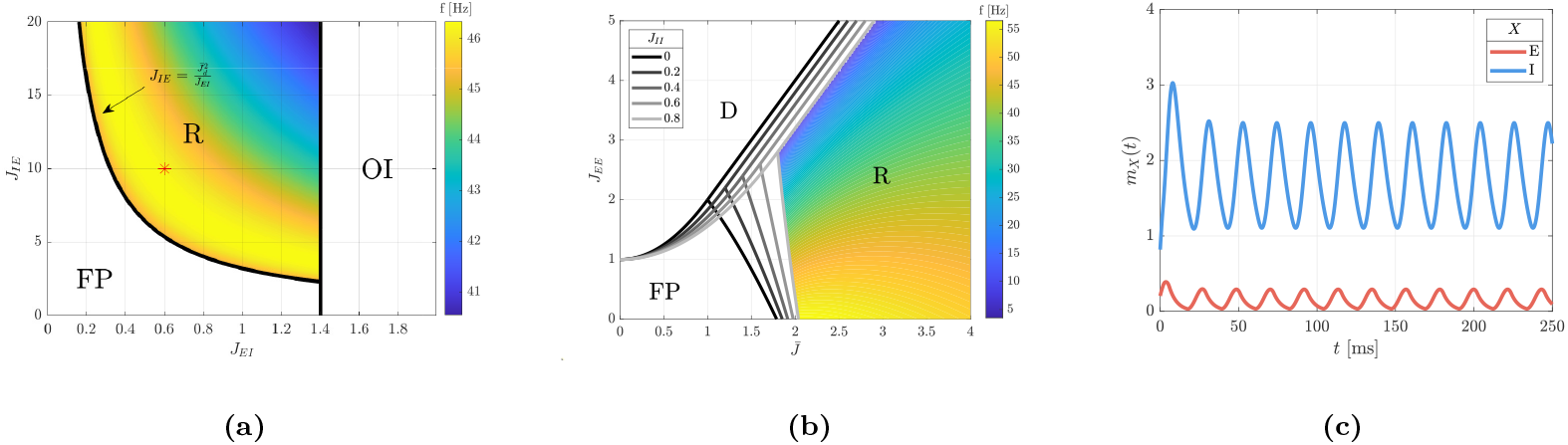
The phase diagram. (a) Phase diagram in the plane of [*J*_*EI*_, *J*_*IE*_]. Three regions are delineated: Fixed Point (FP), Only Inhibition (OI), and Rhythmic (R). The bifurcation line separating the FP and R regions is given by 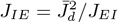. The frequency of the rhythmic activity in the R region is shown by color. Here, we used *J*_*EE*_ = 0.6 and *J*_*II*_ = 0.4. (b) Phase diagram in the plane of 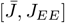. Three regions are delineated: Fixed Point (FP), Rhythmic (R), and Divergence (D) for *J*_*II*_ = 0, 0.2, …, 0.8. The frequency of the rhythmic activity in the R region is shown by color. (c) Example of the mean E and I firing rates in the rhythmic region, depicting *m*_*E*_ (*t*) and *m*_*I*_ (*t*) as a function of time, for *J*_*EE*_ = *J*_*EI*_ = 0.6, *J*_*IE*_ = 10 and *J*_*II*_ = 0.4 (see red asterisk in fig. 2a). For all figures, *τ*_*m*_ = 2.5*d* = 5 ms was used.

When the E-to-E connectivity exceeds a certain thresh-old, 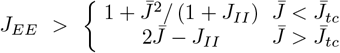,where 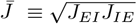,and 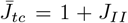 the positive feedback results in a Divergence (D) of the neuronal activities. This scenario represents pathological activity where diverging neuronal responses are suppressed due to a saturating non-linearity that is not described in our model.

For moderate to low I-to-E and E-to-E levels; i.e., outside of the OI and D regions, the system has a Fixed Point (FP) that represents an asynchronous state in which both populations are active,

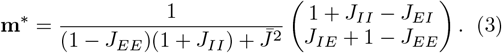

However, when 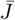 crosses a certain threshold, 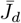, the asynchronous fixed point loses its stability and a limit cycle emerges so that the network relaxes to a rhythmic (R) activity (see figs. 2a and 2c). As the E-to-E connectivity, *J*_*EE*_, is strengthened, the oscillation frequency in the R region decreases monotonically and approaches zero on the transition to the D region, 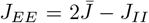(see fig. 2b).

Solving the delayed differential equations for the neuronal dynamics in the R domain is not an easy task.

However, on the bifurcation line the dynamics is linear, yielding:

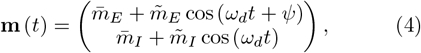

where 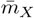 is the temporal mean of the firing rate, 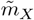 is the amplitude modulation of the firing rate, *ω*_*d*_ is the angular frequency and *ψ* is the relative temporal phase between the excitatory and inhibitory populations (see the derivations for *ω*_*d*_ in Appendix A 1 and for 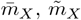 and *ψ* in Appendix A 2).

### B. Beyond order parameters

In the following section we investigate STDP dynamics in which the synaptic weights change over time. The driving force of the STDP dynamics is the product of the STDP rule and the neuronal cross-correlations. This makes it imperative to understand how the crosscorrelations depend on the synaptic weights. To this end, we describe the weights by their means, which serve as the order parameters, and the fluctuations around the means:

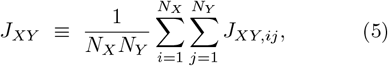

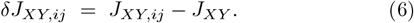

In our model, the cross correlations between different neurons are given by the product of their mean firing rates. We define the temporal averaged cross correlations Γ_*XY,ij*_ (Δ) = ⟨*m*_*X,i*_ (*t*) *m*_*Y,j*_ (*t* + Δ)⟩ _*T*_, where ⟨·⟩_*T*_ denotes averaging over one period of the limit cycle (in the R region). Expanding to first order in *δ***J**, we write

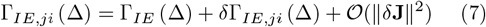

In the FP region, in which the firing rates converge to a fixed point, to a leading order in the fluctuations, the cross correlations are given by (see Appendix A 3 and A 4)

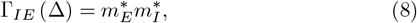

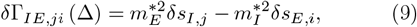

where 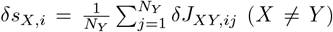 is the mean fluctuation in the inter-synapses to neuron *i* in population *X*.

In general, in the rhythmic region the dynamics is nonlinear and the computation of the limit cycle is not trivial. However, near the bifurcation line we can approximate the neural responses by their dynamics on the bifurcation line, which is linear, yielding (see Appendix A 3 and A 4)

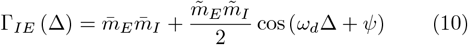

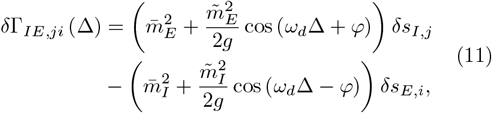

where 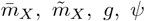 and *φ* are given in Appendix A 1 and A 2. Note that Γ_*EI,ij*_ (Δ) = Γ_*IE,ji*_ (− Δ). For convenience, we further define the following quantities dependent on the order parameters

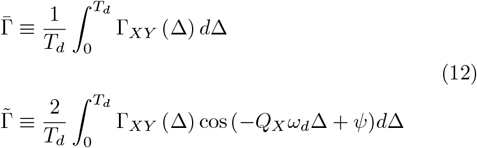

where *T*_*d*_ = 2*π/ω*_*d*_, *Q*_*E*_ = 1, and *Q*_*I*_ = −1. In the FP region 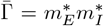 and 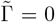. In the R region near the bifurcation line 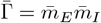 and 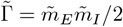.

## III. STDP DYNAMICS

In section III A we define the STDP learning rule and introduce the STDP dynamics in the limit of slow learning. Next, in section III B we study the induced STDP dynamics of the order parameters. Then, in section III C we study the dynamics of the fluctuations around the order parameters.

### A. The STDP rule

We now define the synaptic plasticity rule. Since the inter-connections, *J*_*EI*_ and *J*_*IE*_, are the essential synaptic connections needed to generate rhythmic activity, we focus on their plasticity, and assume that the intraconnections, *J*_*EE*_ and *J*_*II*_, are not plastic (illustrated in fig. 1). In our STDP learning rule, the modification of the weight, *dJ*_*XY,ij*_, of a synapse from pre-synaptic neuron *j* in population *Y* to post-synaptic neuron *i* in population *X*, resulting from a pre-synaptic spike at time *t*_*j*_ and a post-synaptic spike at time *t*_*i*_ = *t*_*j*_ + Δ is given by (see also [36, 43, 45])

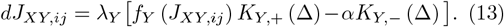

The change in the synaptic weight *dJ*_*XY,ij*_ is given by the contribution of two processes: potentiation (+) and depression (−). Each process can depend on both the spike time difference, Δ, and on the synaptic weight, *J*_*XY,ij*_. Here for mathematical convenience we assumed a separation of variables, and wrote each process as the product of the synaptic weight-dependent function, *f*_*Y*_ (*J*_*XY,ij*_), and the temporal kernel *K*_*Y,±*_ (Δ). The parameters *λ*_*Y*_ and *α* correspond to the learning rate and the relative strength of depression, respectively. Note that since only two types of synapses exhibit plasticity, it suffices to denote *λ*_*Y*_, *f*_*Y*_ and *K*_*Y,±*_ solely by the type of their presynaptic neuron, *Y*.

The weight-dependent function, *f*_*Y*_, restrains the potentiation of Y-to-X synapses from diverging. Following Gütig et al. [36] (see also [43]) for the excitatory synapses we chose

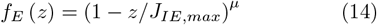

where *µ* characterizes the non-linearity of the weight dependence and *J*_*IE,max*_ is the maximal value of the excitatory synaptic weights. For inhibitory synapses we took *f*_*I*_ = 1.

For the temporal kernels of the STDP rule we focused on temporally asymmetric Hebbian [34, 38, 41, 46, 47] and anti-Hebbian rules [37, 45, 48]. Specifically, we used exponential kernels:

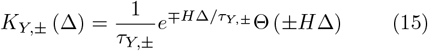

where *τ*_*Y*,+_ and *τ*_*Y,−*_ denote the characteristic time constants for potentiation and depression, respectively. Following [34, 39, 49], we chose *τ*_*Y*,+_ *< τ*_*Y,−*_, see Appendix A 11 for an explanation about the choice of parameters.

The change in the synaptic weight, eq. (13), during short time intervals, yields (see also [36, 45, 50])

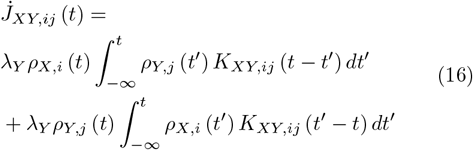

where *K*_*XY,ij*_ (*z*) = *f*_*Y*_ (*J*_*XY,ij*_) *K*_*Y*,+_ (*z*) − *αK*_*Y,−*_ (*z*). In the limit of slow learning, *λ*_*Y*_ ≪ *ω*_*d*_, the STDP dynamics, eq. (16), averages the neural responses over time, yielding (see [45] for a more detailed derivation)

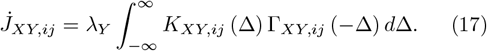

### B. Induced dynamics of the order parameters

Assuming a uniform solution in which *J*_*XY,ij*_ = *J*_*XY*_ reduces the complex high-dimensional dynamics in eq. (17) to the following synaptic dynamics

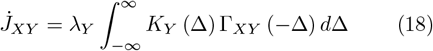

where *K*_*Y*_ (Δ) = *f*_*Y*_ (*J*_*XY*_) *K*_*Y*,+_ (Δ) −*αK*_*Y,−*_ (Δ). The separation of time scales between neural and STDP dynamics enables examination of the neural activity while maintaining constant synaptic weights. In this framework, the synaptic weights *J*_*XY*_ are treated as parameters of the firing rates *m*_*X*_ (*t*) phase diagram (see figs. 2a and 2b). According to eq. (18), on a large time scale, *λ*^*−*1^, STDP induces a dynamic flow on the phase diagram, simultaneously defined as the phase space of synapses [*J*_*EI*_, *J*_*IE*_]. The STDP parameters shape this flow and indirectly dictate the collective behavior of the neural network *m*_*X*_ (*t*) in accordance with the current position of the synaptic weights on the phase diagram.

Substituting Γ_*XY*_ from eqs. (8) and (10) into eq. (18) yields

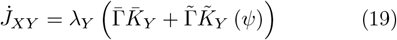

Where

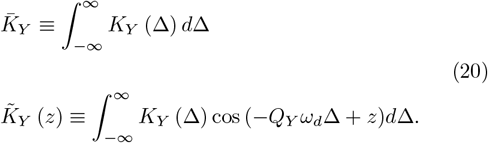

The flow on the phase diagram, eq. (19), is controlled by the first two temporal Fourier transform components of the mean cross-correlations between populations and the learning rule. In the FP region only the first term exists, 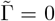 therefore, due to our choice of normalizaTion 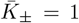, we have 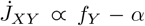. Consequently, for a weak relative strength of depression characterized by *α < f*_*E*_ *<* 1, STDP induces a flow that attracts the system from the majority of the FP region toward the R region, as illustrated in fig. 3a. When the order parameters traverse the bifurcation line into the R region, 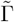 undergoes a discontinuous change from 0 to 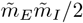. Therefore, as the second term in eq. (19) begins to influence the flow on the phase diagram, it may lead to a change in the sign of 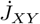. The effect of the second term in eq. (19) on the dynamics will be also determined by the second Fourier transform component of the temporal kernel,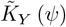. In the case of an asymmetric Hebbian rule for *J*_*EI*_ and an asymmetric anti-Hebbian rule for *J*_*IE*_ we get

**FIG. 3:**
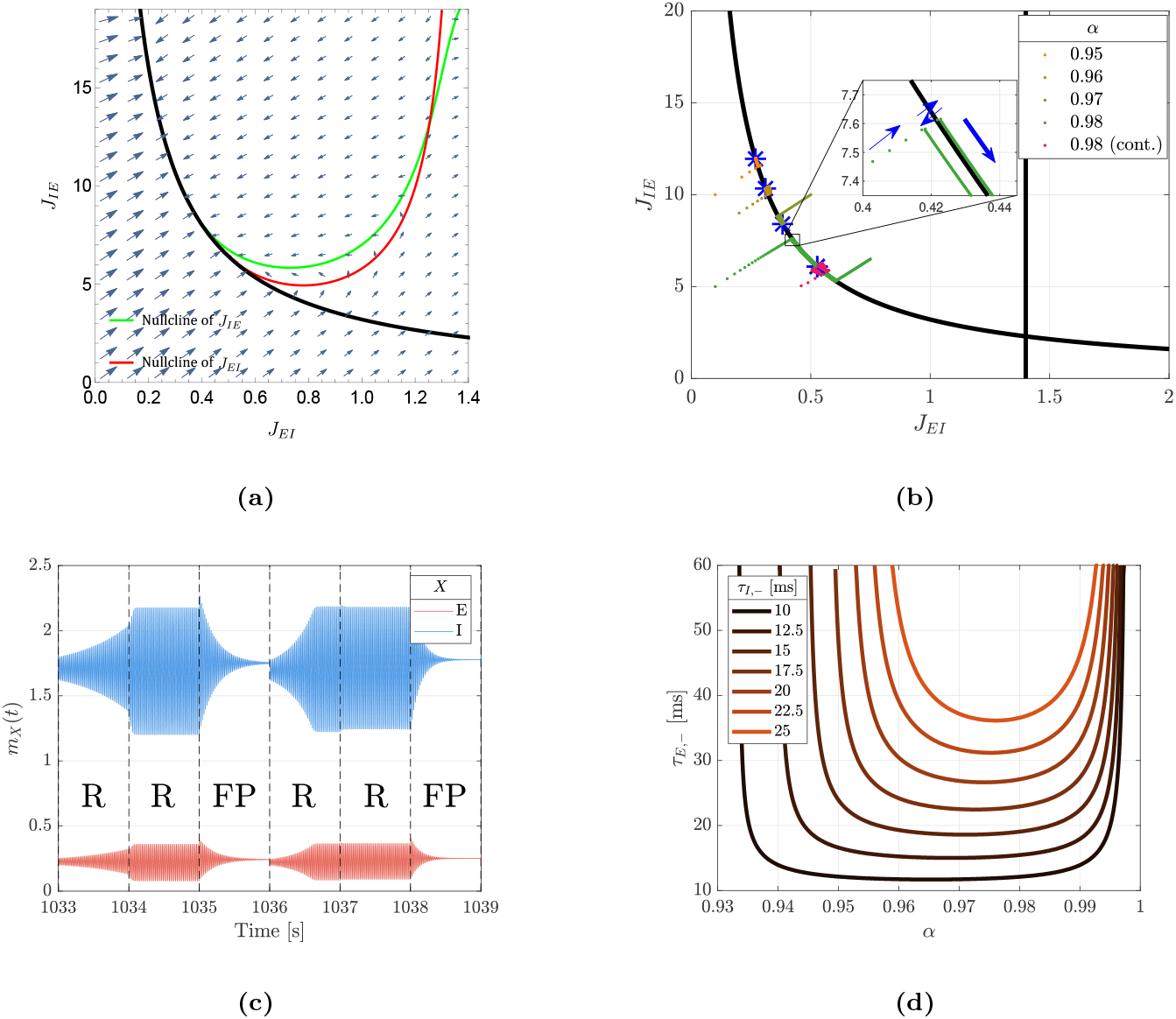
Critical Rhythmogenesis. (a) STDP induces a flow on the phase diagram of the system. The flow is depicted by arrows on the plane of [*J*_*EI*_, *J*_*IE*_]. The black line denotes the bifurcation line. The green and red lines denote the nullclines of the excitatory (*J*_*IE*_) and inhibitory (*J*_*EI*_) mean synaptic weights (order parameters), respectively. Here we used: *α* = 0.96, *τ*_*E,−*_ = 35 ms and *λ*_*E*_ = 10*λ*_*I*_ = 1. (b) STDP dynamics of the order parameters, eq. (18), were simulated for different values of *α* = 0.95, 0.96, 0.97, 0.98 (orange to green), using the separation of timescales (see Appendix A 9). For *α* = 0.98 (green), the dynamics were computed for two different initial conditions. Critical points **J** = **J**^*∗*^ for each *α* are marked with blue asterisks. The pink traces depict the STDP dynamics using continuous rates, for *α* = 0.98 and *λ*_*E*_ = 10*λ*_*I*_ = 3 (see details in Appendix A 9). (Inset) The thin blue arrows highlight the arrival to the bifurcation line and the alternate transitions across it. The thick blue arrow highlights the drift along the bifurcation line towards **J**^*∗*^. Here we used *τ*_*E,−*_ = 25 ms and *λ*_*E*_ = 10*λ*_*I*_ = 4. (c) The mean firing rates of the excitatory (red) and inhibitory (blue) populations for STDP dynamics with continuous rates (pink traces in fig. 3b) when the network is in critical rhythmogenesis. The synaptic weights are updated every 1 s, and ‘FP’ and ‘R’ mark the durations in which the system is in the FP and R regions, respectively. (d) The region in parameter space–specifically here the plane of [*α, τ*_*E,−*_]–in which critical rhythmogenesis can occur, is shown for various values of *τ*_*I,−*_ = 10, 12.5, …, 25 ms. Critical rhythmogenesis is achievable above the corresponding lines. In all the figures we used *µ* = 0.01, *J*_*IE,max*_ = 20, *τ*_*E*,+_ = *τ*_*I*,+_ = 10 ms, *τ*_*I,−*_ = 15 ms, *J*_*EE*_ = 0.6, *J*_*II*_ = 0.4 and *τ*_*m*_ = 2.5*d* = 5 ms.

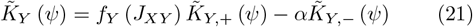

where 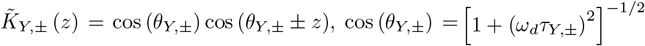. Due to *θ* and *ψ*, the term 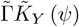 is negative, resulting in a sign change of 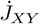 across a significant part of the bifurcation line, as depicted in fig. 3a. Usually a change of sign in the time derivative of all the dynamic variables may indicate an attractor, where the dynamics slowly converge towards it. However, in this case, due to the discontinuity in 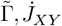 change their sign discontinuously, resulting in repetitive transitions between the FP and R regions (inset in fig. 3b). These alternating transitions generate a continual toggling between fixed firing rates and rhythmic activity. Figure 3c depicts the mean firing rates of the excitatory and inhibitory populations during these alternating transitions. In this simulation the synaptic weights are updated every 1s, marked by the dashed black lines. ‘R’ and ‘FP’ denote whether the system is in the R or FP regions. Note that the firing rate dynamics exhibits a transient behavior. A qualitatively similar behavior has been described in [43] in a reduced model only with inter-connections.

The transitions between the FP and R regions exhibit a drift along the bifurcation line, as shown in the inset of fig. 3b. This drift arises because the direction of 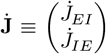 on one side of the bifurcation line does not necessarily oppose the direction of 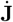 on the subsequent step in the other region. When such opposition occurs at a specific point on the bifurcation line, denoted **J**^*∗*^, the drift vanishes and the system settles into repetitive transitions around this critical point (see derivation of **J**^*∗*^ in Appendix A 5).

In fig. 3b, the dynamics of the order parameters are illustrated for varying values of the relative strength of depression *α*. For each *α* the system achieves critical rhythmogenesis at different values of **J**^*∗*^ (indicated by the blue asterisks). Figure 3d illustrates the phase diagram for critical rhythmogenesis by delineating its domain within the [*α, τ*_*E,−*_] plane (see details on the conditions for critical rhythmogenesis in Appendix A 5). Each line represents the boundary for different *τ*_*I,−*_ values, above which critical rhythmogenesis emerges.

### C. STDP dynamics of the fluctuations around the order parameters

We next examined the STDP dynamics with synaptic fluctuations around the uniform solution, while the order parameters are relaxed to critical rhythmogenesis. Assuming small fluctuations, we expand eq. (17) to the leading order in the fluctuations, *δJ*_*XY,ij*_, around **J**^*∗*^, to obtain

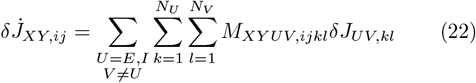

where *M*_*XY UV,ijkl*_ is a tensor relating a fluctuation in synaptic weight *δJ*_*UV,kl*_ to the dynamics of a fluctuation in the synaptic weight *δJ*_*XY,ij*_. The tensor depends on the order parameters **J**^*∗*^, and on the collective state of the system; i.e., FP or R states. The details for *M*_*XY UV,ijkl*_ appear in Appendix A 6.

The eigenvectors of the tensor *M*_*XY UV,ijkl*_ can be classified into four families which represent four qualitatively different structures in which synaptic fluctuations may evolve in time. These types of synaptic fluctuations are orthogonal to the uniform configuration of the synapses. Moreover, these families emerge regardless of the network size (although not for *N*_*E*_ = 1 and/or *N*_*I*_ = 1), so that we simply visualized them for *N*_*E*_ = 3 and *N*_*I*_ = 2 in fig. 4.

**FIG. 4:**
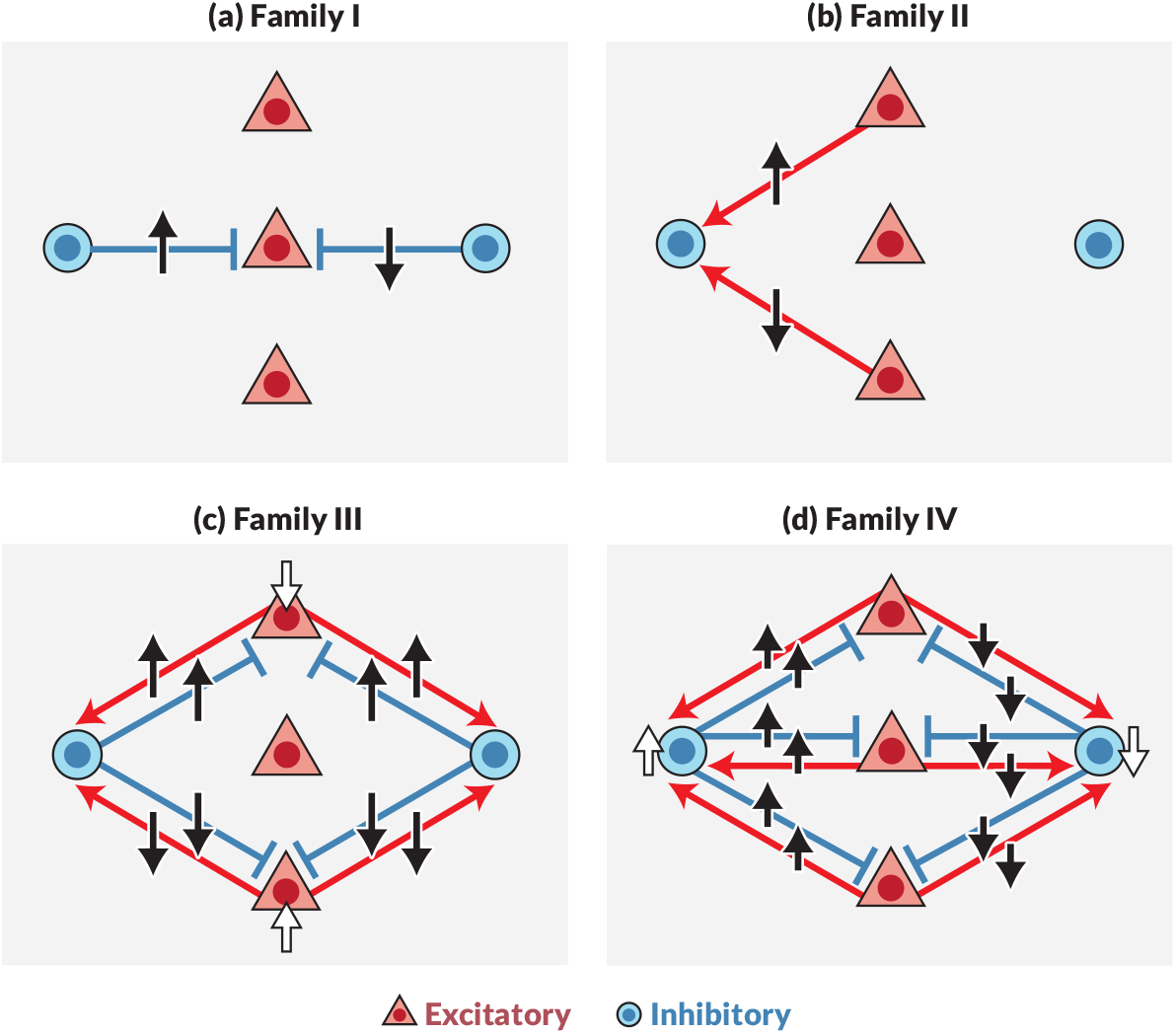
Illustrations of the four families of synaptic fluctuations, for *N*_*E*_ = 3 and *N*_*I*_ = 2. (a) Family I fluctuations in the I-to-E synapses, such that the inhibitory input to each excitatory neuron remains fixed. (b) Family II - fluctuations in the E-to-I synapses, such that the excitatory input to each inhibitory neuron remains fixed. (c) Family III - fluctuations in which excitatory neurons change their activity, while keeping the mean excitatory activity fixed. (d) Family IV - fluctuations in which inhibitory neurons change their activity, while keeping the mean inhibitory activity fixed. Upright (inverted) black arrows indicate strengthening (weakening) of the synapse. The white arrows illustrate the change in neural activity.

Eigenvectors in family I (fig. 4a) are fluctuations in inhibitory synapses to an excitatory cell, *k* (*k* ∈ {1, … *N*_*E*_}), such that ∑*l* Δ*J*_*EI,kl*_ = 0 and thus conserve the total inhibitory input to neuron *k* and consequently do not affect the response dynamics. The constancy in neural dynamics does not alter the STDP dynamics, resulting in marginal stability for any order in these synaptic fluctuations. The multiplicity of these eigenvectors is *N*_*E*_ (*N*_*I*_ −1).

Eigenvectors in family II (fig. 4b) are fluctuations in the excitatory synapse s to an inhibitory cell, *k* (*k* ∈ {1, … *N*_*I*_}), such that ∑*l* Δ*J*_*IE,kl*_ = 0 and thus conserve the total excitatory input to neuron *k* and consequently do not affect the response dynamics. Although constancy in neural dynamics does not alter the STDP dynamics, built-in depression in excitatory dynamics, due to *f*_*E*_, suppresses these fluctuations, resulting in a negative (stable) eigenvalue (see Appendix A 6). The multiplicity of these eigenvectors is *N*_*I*_ (*N*_*E*_ − 1).

In family III (fig. 4c) the firing rates of individual excitatory neurons change while keeping the mean excitatory activity, *m*_*E*_(*t*) = *i m*_*E,i*_(*t*), as well as activity of the inhibitory neurons fixed. In the example in fig. 4c, the white arrows indicate the neurons that change their activity. In the eigenvectors in family III, all the inhibitory connections to excitatory neuron *k* are modified by the same amount, 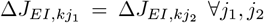, and all of the connections of excitatory neuron *k* to the inhibitory neurons obey 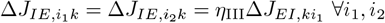, *i*_2_ (see Appendix A 6). Figure 4c depicts an example eigenvector for this family. In this example, the fluctuations in the excitatory firing rates are in the form of *δm*_*E*_ = (− 1, 0, 1) (from top to bottom). This is achieved by increasing all the inhibitory connections to the top excitatory neuron and decreasing all the inhibitory connections to the bottom neuron. Fluctuations in this family decay (negative eigenvalue) when the network is around the bifurcation line. The degeneracy of this eigenvector results from the constraint *i m*_*E,i*_ = *const*; hence, this family spans an *N*_*E*_ − 1 dimensional space.

In family IV (fig. 4d), firing rates of individual inhibitory neurons change while keeping the mean inhibition, *m*_*I*_ (*t*) = *j m*_*I,j*_(*t*), fixed. In these eigenvectors, all the excitatory connections to inhibitory neuron *k* are modified by the same amount, 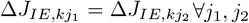, and all of the connections of inhibitory neuron *k* to excitatory neurons obey 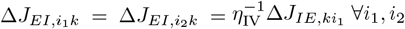 (see Appendix A 6). Family IV eigenvectors span an *N*_*I*_ − 1 dimensional space from the condition ∑ *j m*_*I,j*_ = *const*.

The stability analysis of fluctuations in this family is more intricate. We found that the eigenvalue corresponding to this family is typically positive (unstable) in the FP region, 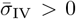, and negative (stable) in the R region, 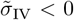, as shown in fig. 5a. Thus, as the system switches alternately from FP to R states around the critical point, **J**^*∗*^, these fluctuations grow (in the FP) and then decay (R) repeatedly. What can be said about the net effect on these fluctuations?

**FIG. 5:**
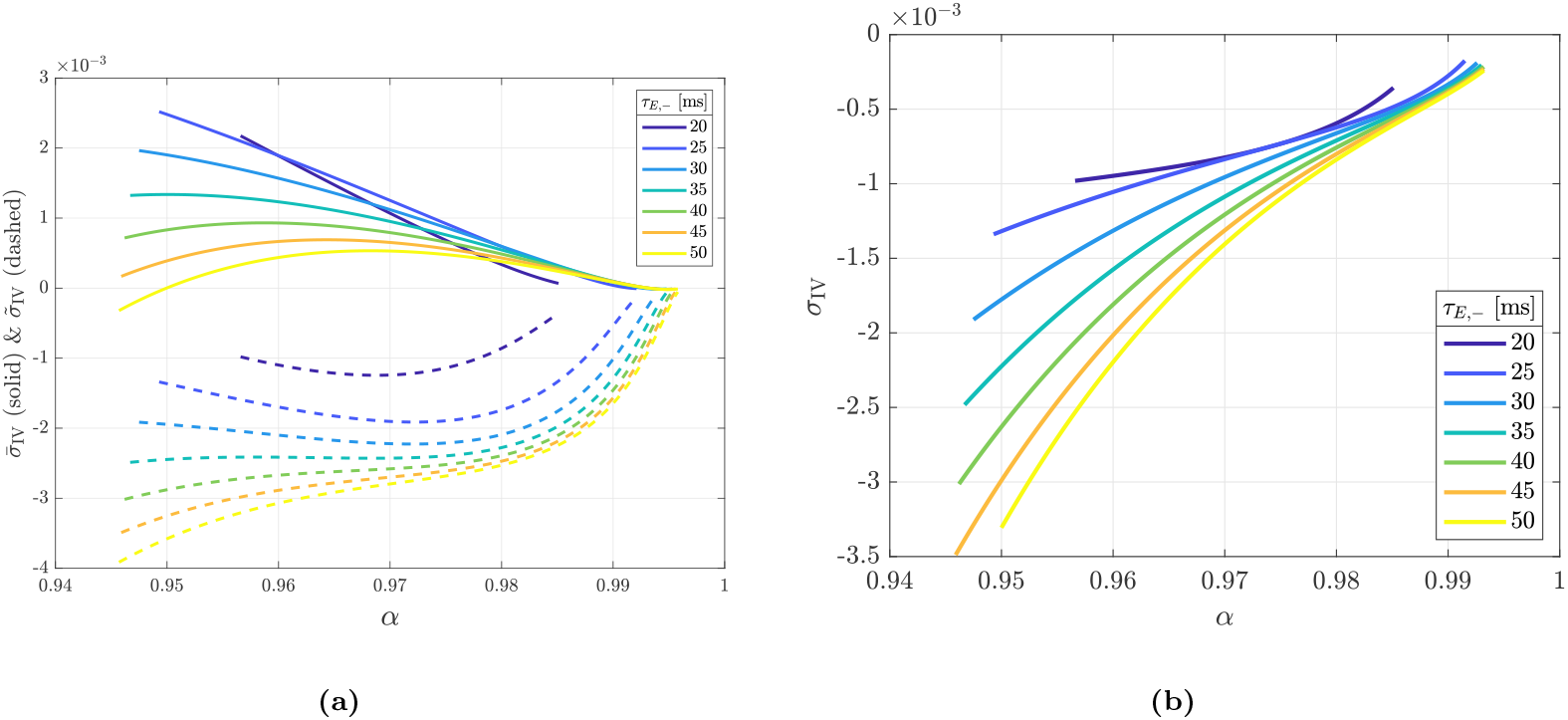
The eigenvalues of family IV. (a) The eigenvalues of family IV in the FP regime (solid) and R regime (dashed) near the bifurcation line. (b) The value of *σ*_IV_, which governs the stability of family IV fluctuations in critical rhythmogenesis. Each figure is drawn as a function of *α* for different values of *τ*_*E,−*_ = 20, 25, …, 50 ms differentiated by color. We used *µ* = 0.01, *J*_*IE,max*_ = 20, *λ*_*E*_ = *λ*_*I*_ = 1, *τ*_*E*,+_ = *τ*_*I*,+_ = 10 ms, *τ*_*I,−*_ = 15 ms, *J*_*EE*_ = 0.6, *J*_*II*_ = 0.4 and *τ*_*m*_ = 2.5*d* = 5 ms.

Heuristically, consider the following scenario. During the time, 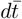, the system spends in the FP state, the fluctuations grow by a factor 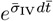. During the time, 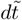, the system spends in the R state, the fluctuations decay by a factor 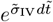. Thus, stability is governed by the effective stability eigenvalue of family IV, defined as the weighted mean of the eigenvalues, with weights given by the relative duration spent in each state:

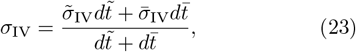

see Appendix A 7). The duration spent in each region can be estimated by the inverse of the absolute value of the time derivative of the order parameters, 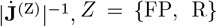. At the critical point,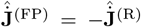; therefore, the magnitude of the ratio of 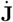 is equal to the ratio of the magnitude of each of its components, 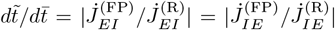. Figure 5b depicts the effective eigenvalue of family IV, *σ*_IV_, for different values of *α* and *τ*_*E,−*_. As shown, the effective eigenvalue is negative and fluctuations in this family of eigenvectors are expected to decay over time.

To study the stability of critical rhythmogenesis with respect to fluctuations, we simulated STDP dynamics with additive Gaussian white noise (see details on white noise statistics in Appendix A 9). The magnitude of the fluctuations in each family as a function of time is depicted in fig. 6a. As can be seen in the figure, the fluctuations in families II, III and IV converged to a steady state. For comparison, the dashed line depicts the asymptotic variance of family IV, as computed analytically assuming small fluctuations, see Appendix A 7. These fluctuations are correlated in time over a timescale of 1*/* |*σ*_IV_| ≈ 160 [a.u.], fig. 6b (see details of the analytical derivation in Appendix A 7). By contrast, fluctuations in family I, with marginal stability, accumulated over time. Nevertheless these fluctuations did not affect the neuronal response dynamics. This can be seen in figs. 6c and 6d, which show that the variability of the neural responses across the population converged to a finite size.

**FIG. 6:**
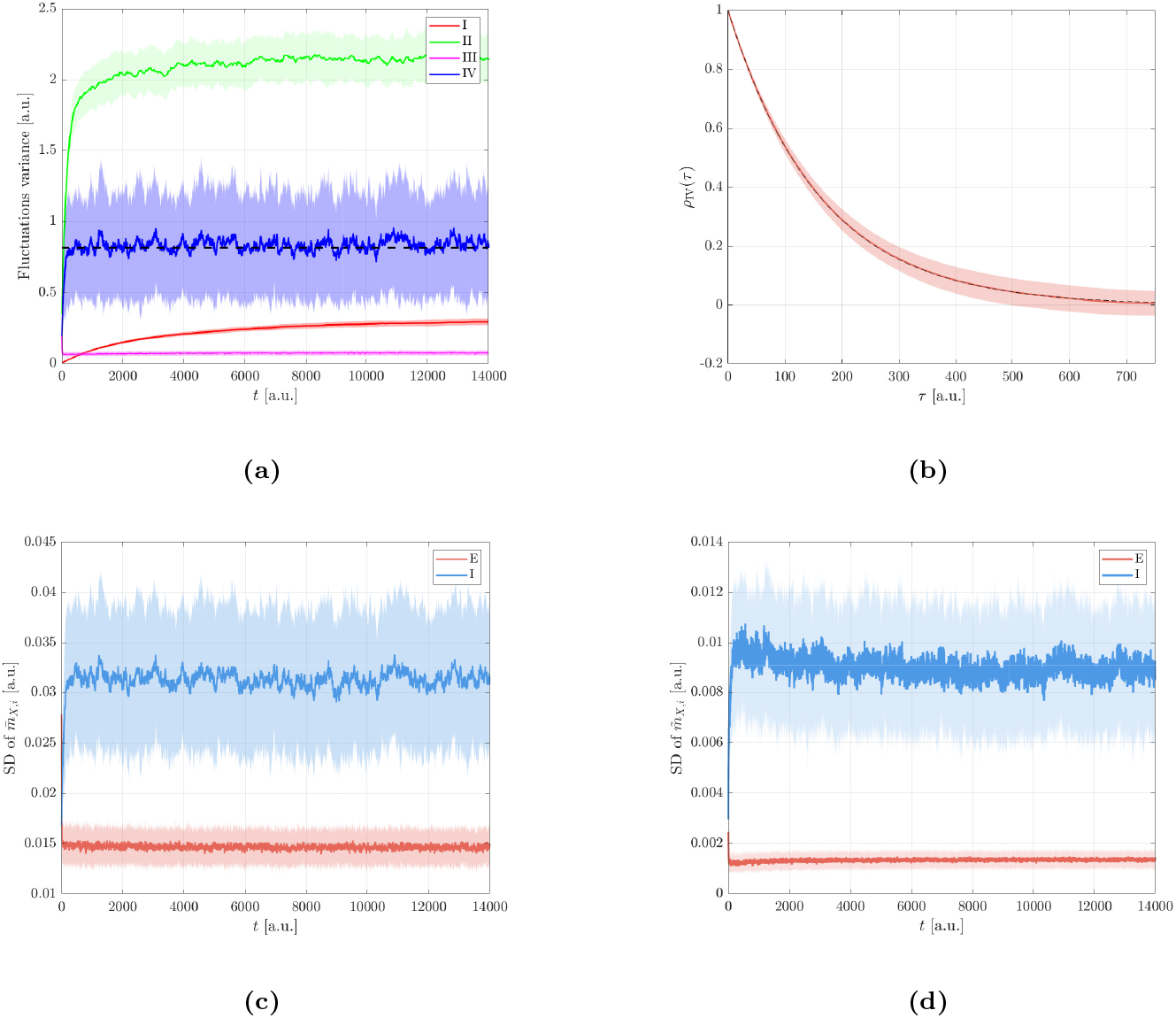
Noisy STDP dynamics. STDP dynamics were simulated with additive Gaussian white noise (see details in Appendix A 9). Results were averaged over 100 noise realizations; the solid lines represent the means, and the shaded regions indicate the standard deviations. The initial conditions for the synaptic weights were drawn independently from Gaussian distributions with *J*_*XY,ij*_ ∼*N* (*J*_*XY*,0_, [*J*_*XY*,0_*/*10]^2^), with *J*_*EI*,0_ = 0.52 and *J*_*IE*,0_ = 6. After a short transient, the system converges to critical rhythmogenesis. (a) The fluctuations in each family (differentiated by color) are shown as a function of time. At each time step, the fluctuations were decomposed into the sum of the four vectors, one from each family. For each family, we plot the squared Euclidean norm per dimension, ∥**x**∥ ^2^*/*dim. The dashed black line presents the asymptotic variance in the subspace of family IV (see Appendix A 7). (b) Normalized autocorrelation of family IV. The correlation coefficient of fluctuations in family IV at different times (in equilibrium), *ρ*_IV_(*τ*), is shown as a function of the time difference, *τ*. The solid red line and shaded regions depict numerical estimation. The dashed black line shows analytical result; see details in Appendix A 7. (c) The standard deviation in 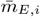 (red) and 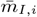 (blue) across their respective populations as a function of time. (d) The standard deviation in 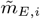 (red) and 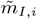 (blue) across their population as a function of time. The neural and STDP dynamics (eqs. (1) and (17)) were simulated for a network consisting of *N*_*E*_ = 40 excitatory and *N*_*I*_ = 10 inhibitory neurons. Shaded areas represent the standard deviation across 100 trials. Here we used *α* = 0.98, *µ* = 0.01, *J*_*IE,max*_ = 20, *λ*_*E*_ = 10*λ*_*I*_ = 10, *τ*_*E*,+_ = *τ*_*I*,+_ = 10 ms, *τ*_*E,−*_ = 25 ms, *τ*_*I,−*_ = 15 ms, *J*_*EE*_ = 0.6, *J*_*II*_ = 0.4 and *τ*_*m*_ = 2.5*d* = 5 ms.

## IV. DISCUSSION

Modifications of synaptic weights are considered one of the fundamental mechanisms underlying learning in the brain. STDP provides a microscopic learning rule operating at the synaptic level. Because STDP operates independently of external reward or error signals, it is typically regarded as an unsupervised learning rule. In the absence of such supervisory signals, unsupervised learning is often guided by an intrinsic computational principle—for example, dimensionality reduction, clustering, or homeostasis. Thus, raising the question: what possible computational principles can STDP implement? Here, we investigated whether STDP can serve as a mechanism for the emergence of temporal patterns in neuronal activity.

Here we extended previous research on critical rhythmogenesis [43] (see also [48]). Our approach to investigating recurrent STDP dynamics has two stages. First, the dimensionality of the problem is reduced by studying the induced STDP dynamics of the order parameters. Then, a perturbative analysis is carried out in the subspace that is orthogonal to the order parameters.

The interconnections; i.e., E-to-I and I-to-E, are essential for generating the rhythmic activity in the specific model studied here [23, 43]. The intra-connections (E-to-E and I-to-I) mainly modulate this activity. However, beyond certain critical values, these intra-connections can drive the network toward either a pathological state (fig. 2b) or non-physiological rapid oscillations (see Appendix A 1). We found that homeostatic plasticity of the intra-connections can prevent the drift into these pathological regions and maintain the intra-connections in the desired working range (see Appendix A 8). However, since we were interested in rhythmogenesis, we focused here on the plasticity of the inter-connections and assumed that the intra-connections were non-plastic.

The order parameters span the phase diagram of the neural response (figs. 2a and 2b). Due to the separation of timescales between the neural response and the learning process, the STDP dynamics of the order parameters can be viewed as a flow on the phase diagram. We found that for a wide range of parameters, STDP induced a flow toward the bifurcation line (figs. 3a and 3b). Note that in this case, the system does not converge to a fixed point, and the temporal derivative of the order parameters, 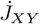, does not vanish on the bifurcation line.

Rather, 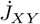 can be positive to the left of the bifurcation line and negative to the right of it, for example. Thus, the attraction to the bifurcation line results from two key features. The first is that STDP is an *activity*-dependent plasticity rule. The second is that activity changes sharply on the bifurcation line. Hence, this mechanism can be generalized to other scenarios. Reducing the dimensionality by focusing on the effective dynamics of the order parameters allowed us to study the effects of the non-linearity of the neuronal responses; i.e., the transition to rhythmic activity. To study the stability of the order parameters to arbitrary fluctuations, a stability analysis was carried out in the orthogonal direction. Our analysis revealed four families of eigenmodes of fluctuations. Two families (II and III) are characterized by negative eigenvalues, indicating decay. One family is characterized by a positive eigenvalue in the FP region and a negative eigenvalue in the R region but the net effect is a decay in this subspace as well.

Finally, one family (I) is characterized by zero eigenvalues. Because it does not affect neural activity, the STDP dynamics of its fluctuations neither grow nor decay,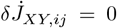. Consequently, this family defines a manifold spanned by its eigenvectors, permitting the system to move freely within it (see also Burkitt et al. [51]). Thus, in our model, STDP maintains functionality by stabilizing the order parameters that govern the function while allowing considerable fluctuations in an orthogonal subspace.

In this study, we examined functionality in terms of rhythmogenesis—the network’s capacity to autonomously generate rhythmic activity [52]. Across a wide range of parameters, the dynamics induced by STDP do not converge to a stable fixed point within the rhythmic region (as reported in [52, 53]) but instead settle at the boundary between rhythmic and non-rhythmic regimes (see also [43] for discussion about the generic nature of critical rhythmogenesis in this model and about alternative scenarios). Consequently, the system persistently transitions between these two modes of activity. We refer to this operating regime as critical rhythmogenesis, which provides several potential functional advantages.

First, critical rhythmogenesis enables rapid switching between rhythmic and non-rhythmic states—for, for instance through neuromodulatory control, which offers a wider repertoire of dynamic patters of neuronal activities. Second, it emerges robustly over a wide parameter range and is relatively insensitive to specific features of the STDP rule governing interconnections. Finally, activity near criticality has been proposed to offer computational benefits in the brain [54–58].

The question of maintaining functional stability in the face of continuous synaptic motility has been addressed in the past, mainly in the framework of stabilizing a certain spatial structure of the neural responses. Three main approaches have been suggested. One approach assumes that only a subgroup of synapses is essential for functionality, and that these synapses are not (or less) volatile, either due to their type or strength [2, 59–62]. Another approach builds on the observation that neurons can have multiple synaptic connections. In this case, the noise in the effective synaptic weight can be decreased [63–65].

An alternative idea posits that functionality is stabilized through activity-dependent plasticity. Thus, fluctuations in the synaptic connectivity that disrupt functionality will cause a change in the neural activity, which in turn, will induce negative feedback by the activity dependent plasticity. Thus, in this approach, activity dependent plasticity functions as a counterbalancing force that mitigates intrinsic synaptic fluctuations [29–31]. In Manz et al. [28], for example, the target functionality was the formation of a spatial structure in the form of neuronal clusters. Two types of fluctuations were found. The first were fluctuations that could disrupt functionality. These fluctuations were suppressed by STDP. The second were fluctuations that did not disrupt functionality, such as transferring a neuron from one cluster to another. These fluctuations were not suppressed. Thus, STDP allowed the system to drift in the space of possible solutions for network functionality.

Interestingly, Susman and colleagues [66] investigated homeostatic mechanisms that can mitigate the accumulation of random fluctuations in synaptic weights. They found that these mechanisms also degrade the stored memory in the system. However, they reported that memories that were encoded as dynamical patterns were more robust than memories that were encoded by fixed points.

In our work, the target function is the formation of a temporal structure. Here as well, we found two types of fluctuations. Fluctuations of the order parameters, as well as families III and IV may disrupt functionality and are suppressed by STDP. On the other hand, fluctuations in family I do not affect functionality. Hence, STDP allows the system to drift on the manifold of solutions spanned by family I.

At first glance, the analysis of individual synaptic dynamics may appear to simply validate the order parameter framework, suggesting that synaptic fluctuations play a minimal role—consistent with the rapid decay of families III and IV in fig. 6a. However, a deeper examination reveals that STDP not only suppresses fluctuations detrimental to network function (families III and IV), but also permits substantial synaptic variability along a manifold of fluctuations that preserve the network’s rhythmic activity (family I). Crucially, this synaptic flexibility does not lead to clustering, which would otherwise compromise global rhythmicity, thus highlighting a form of constrained variability that supports robust functionality.

In the discussion above, STDP was primarily described as a developmental mechanism for rhythmogenesis. However, STDP can also serve a homeostatic role, conferring robustness to perturbations. To illustrate this, we examined the impact of a perturbation in the form of cell death. Figure 7 presents two examples in which, after the system reached critical rhythmogenesis, 30% of the inhibitory neurons were removed. Immediately after their removal, the mean I-to-E synaptic weight remained largely unchanged, but the total inhibitory input to the excitatory population decreased proportionally. As a result: (*i*) the system transitioned out of critical rhythmogenesis into a fixed point (FP), and (*ii*) the bifurcation line corresponding to the reduced inhibitory population shifted rightward (compare the dashed and solid lines). STDP then guided the system back to the critical rhythmogenesis regime, now relative to the new bifurcation line, as shown in the purple trace in Figure 7.

**FIG. 7:**
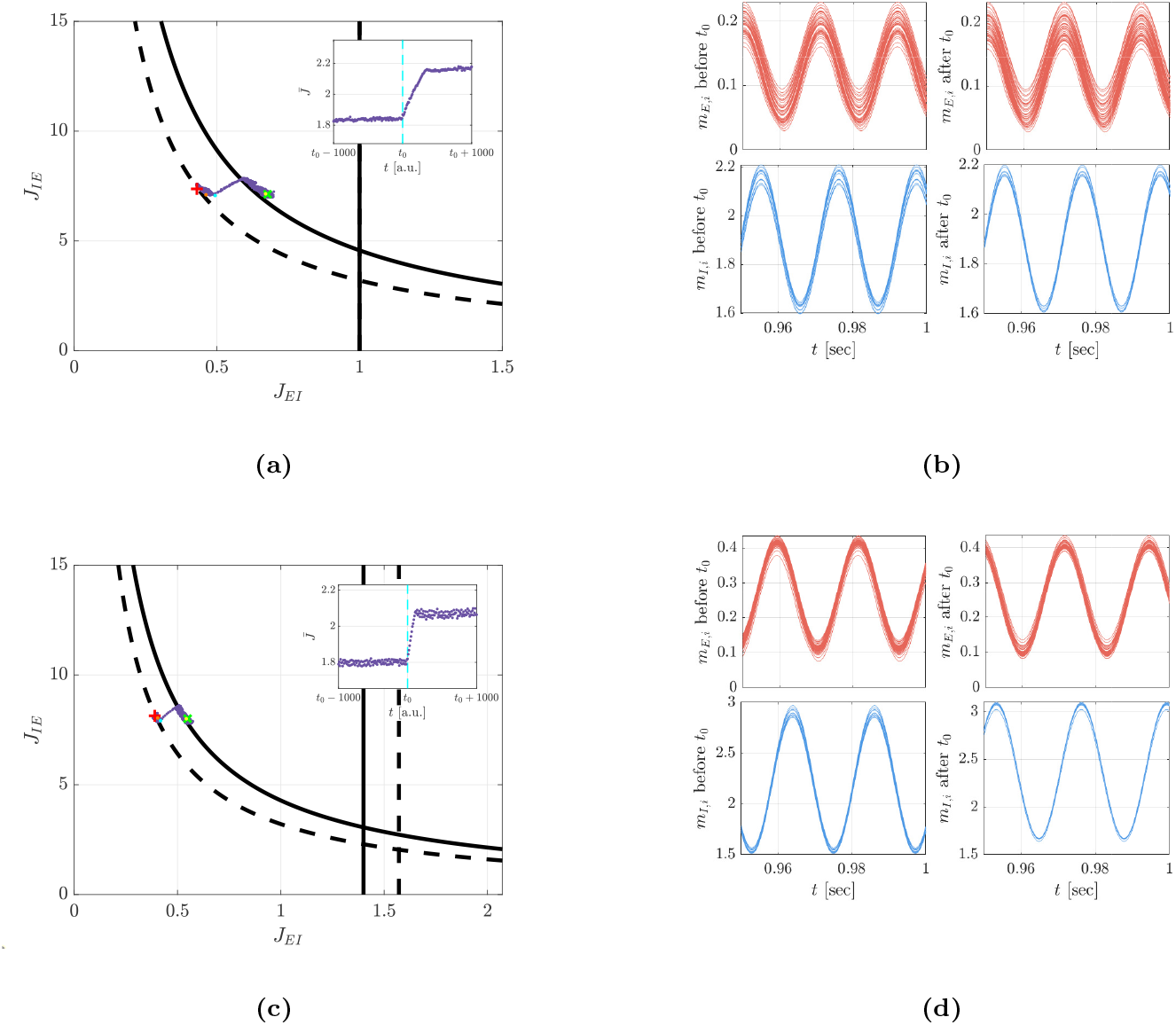
Robustness to cell death. We simulated noisy STDP dynamics of a network with initially *N*_*E*_ = 40 excitatory and *N*_*I*,init_ = 10 inhibitory neurons. At time *t*_0_, three inhibitory neurons were removed (‘killed’): their activity and synaptic connections were set to zero (see Appendix A 9). Importantly, (*i*) the normalization factor *N*_*I*_ in the rate dynamics, eq. (1), remained fixed at *N*_*I*,init_ = 10, and (*ii*) the order parameters (mean synaptic weights) were averaged over the remaining inhibitory neurons, using *N*_*I*_ = *N*_*I*,init_ for *t < t*_0_ and *N*_*I*_ = *N*_*I*,final_ = 7 for *t > t*_0_ in eq. (5). (a) and (c) Trajectories of the order parameters (*J*_*EI*_, *J*_*IE*_) on the phase diagram. Each trajectory begins at the red ‘+’ and ends at the green ‘*×*’, with purple dots marking the temporal evolution. The cyan dot indicates the state at *t*_0_. The dashed and solid black lines depict the bifurcation lines before and after cell death, respectively. Insets show the dynamics of 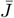 around *t*_0_. The vertical cyan line marks *t*_0_. (b) and (d) show a sample of the mean firing rates in the rhythmic region before (central column, corresponding to the orange dot on the phase diagram in a and c) and after *t*_0_ (right column, corresponding to the yellow dot). Excitatory (*m*_*E,i*_, top red) and inhibitory (*m*_*I,i*_, bottom blue) activities of single neurons are shown as a function of time. In (a) and (b) the intra population connections were set to zero (*J*_*II*_ = *J*_*EE*_ = 0). In (c) and (d), *J*_*EE*_ = 0.6, *J*_*II*_ = 0.4. Initial inter-population weights were drawn from Gaussian distributions: *J*_*EI*_ *∼ N* (0.43, 0.04) and *J*_*IE*_ *∼ N* (7.37, 0.73) in (a), and *J*_*EI*_ *∼ N* (0.39, 0.04) and *J*_*IE*_ *∼ N* (8.14, 0.81) in (b). STDP dynamics, eq. (17), were simulated with Gaussian white noise (see Appendix A 9). Here we used *α* = 0.98, *µ* = 0.01, *J*_*IE*,max_ = 20, *λ*_*E*_= 10*λ*_*I*_ = 10, *τ*_*E*,+_ = *τ*_*I*,+_ = 10 ms, *τ*_*E,−*_ = 35 ms, *τ*_*I,−*_ = 15 ms, and *τ*_*m*_ = 2.5*d* = 5 ms.

Notably, when intra-population connections were absent, figs. 7a and 7b, the frequency of rhythmic activity was very stable (initial and final frequencies: 47.645 ± 0.002 Hz). In contrast, when intra-connections were non-zero, figs. 7c and 7d, the frequency shifted (from 45.0 ± 0.2 Hz to 43.4 ± 0.2 Hz), reflecting changes in inhibitory input to the inhibitory population.

The analysis of STDP in recurrent neuronal networks is challenging due to the associated mathematical difficulties of solving high dimensional coupled non-linear dynamics. The coupling enters the dynamical equations via the cross-correlations of the neuronal responses, which together with the STDP rule provide the driving force for the dynamics. Typically, in order to compute the crosscorrelations, theoretical studies assume that the neural response is linear [28, 51, 67–73]. This assumption does not make the STDP dynamics low dimensional, uncoupled or linear. It simply makes it possible to derive the dynamical equations. To proceed with the analysis, one of two approaches is usually applied.

One approach is to study certain structures, known as motifs [67, 74, 75] analytically and numerically. While this approach is prevalent and addresses specific structures of interest, it fails to provide a comprehensive understanding of the dynamics associated with all synaptic weights.

The other approach reduces the dimensionality by studying the dynamics of the order parameters [28, 51, 67, 69–73]. Then, in some studies, a complementary step is taken that involves performing a linear stability analysis around these order parameters; see e.g. [51].

Here, we applied the latter approach. However, unlike other studies, we did not assume that the neural response was linear. Instead, to facilitate the analysis, we reduced the dimensionality when deriving the cross-correlations of the neural responses. This enabled us to capture the non-linear effects of the neural responses, such as the transition between a fixed-point and a rhythmic activity. How can our theory be tested? The dynamics of STDP are driven by the overlap of neuronal activity and the STDP learning rule. However, experimental preparations that allow direct measurement of the STDP rule typically lack the neuronal activity observed in behaving animals. Conversely, in preparations that enable recording of neuronal activity during behavior, it is nearly impossible to measure the underlying STDP rule. Consequently, deriving empirical predictions for STDP theory is challenging (but see [76] for a notable exception).

Nevertheless, three key predictions emerge from our framework: (1) the system should operate near the boundary between rhythmic and non-rhythmic regimes; Cboth E-to-I and I-to-E synapses should exhibit STDP; and (3) the temporal structure of the STDP rule should be consistent with the families that support rhythmogenesis. For example, under temporally asymmetric rules, the E-to-I STDP kernel should be anti-Hebbian, whereas the I-to-E kernel should be Hebbian.

Our analysis relied on several simplifying assumptions to make the dynamics tractable. In particular, the stability analysis of the STDP dynamics in the R region was facilitated by two key choices. First, we employed STDP rules that lead to critical rhythmogenesis, ensuring that the system remained close to the bifurcation line (for rhythmogenesis via STDP dynamics that lead to a fixed-point in the R region, see [43, 52]). Note also that critical rhythmogenesis has a large basin of attraction in the R region, as is evident by the flow lines in fig. 3a; thus, for long periods of time the system will be near the bifurcation line. Second, we assumed a threshold-linear neuronal response function, eq. (1), which induces a degenerate Hopf bifurcation. This allowed for an exact solution of the neuronal dynamics along the bifurcation line. It is worth noting that we have previously shown that our results can generalize well to systems with smooth neuronal response functions that do not induce a degenerate Hopf bifurcation ([43]; see, for example, Fig. 5c therein). We applied our approach to investigate the emergence of a specific function within a particular model architecture. We did not consider a richer temporal repertoire for the neuronal responses, such as incorporating short term plasticity (but see [52, 53]), burst activity (but see [77]) or others. Regarding the important issues of sparse connectivity and structural plasticity, fig. 9 and Appendix A 10 show that critical rhythmogenesis can tolerate a certain degree of diluted connectivity. These numerical results, however, do not constitute a comprehensive treatment of these fundamental topics, which are beyond the scope of the current study and will be addressed elsewhere. Nevertheless, we believe that our framework can be utilized to explore the development and maintenance of a wide range of dynamic activities in the central nervous system, including the generation of rhythmic patterns by Central Pattern Generators, the sharpening of stimulus selectivity, predictive coding, and memory formation. By capturing the interplay between synaptic plasticity and network dynamics, our approach offers a foundational tool for addressing these complex phenomena.

## Appendix A

**Description of the analytical derivations**

### 1. Stability analysis of the homogeneous fixed-point solutions

The firing rate dynamics for a homogeneous neural activity is given by

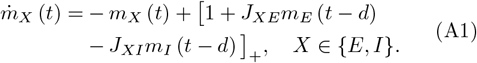

These dynamics have two kinds of fixed point solutions, 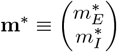. In one (OI) the inhibitory population fully suppresses the excitatory activity; i.e., 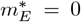, with the requirement that the total input to the excitatory population be negative, 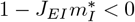, leading to

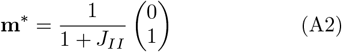

and the existence condition for this solution; i.e., complete suppression of the excitatory population, is *J*_*EI*_ *>* 1 + *J*_*II*_. In order to study the stability of the OI state we introduce a perturbation 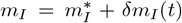, with *δm*_*I*_ (*t*) = *δm*_*I*_*e*^*λt*^. The OI fixed point loses its stability in a Hopf bifurcation, *λ* = *iω* (*ω* ∈ R) for

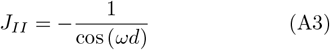

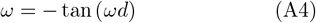

where the solution to these equations is found in *ω*∈ [*π/*2*d, π/d*].

In the case where the excitatory population is not fully suppressed, *J*_*EI*_ *<* 1 + *J*_*II*_, the fixed point (FP) is given by

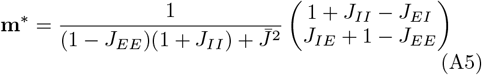

where 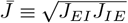

Introducing a perturbation **m**^*∗*^ = **m**+ *δ***m**(*t*), with *δ***m** (*t*) = *δ***m***e*^*λt*^, to this fixed point we obtain from eq. (A1)

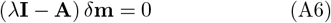

Where

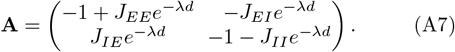

Requiring det [*λ***I** − **A**] = 0 leads to

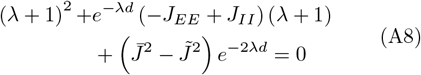

Where 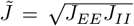. Solving for *λ* + 1 results in two equations for the eigenvalues

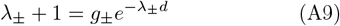

Where

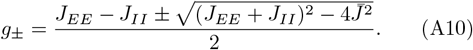

For 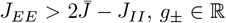. If, in addition, *λ*_*±*_ ∈ ℝ, instability toward the D state occurs when *g*_+_ *>* 1; i.e.,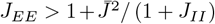. For 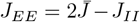, then *g*_+_ = *g*_*−*_ ∈ ℝ, and if *λ*_+_ = *λ*_*−*_ ∈ ℝ, then instability towards the D regime occurs when *g*_*±*_ *>* 1; i.e., *J*_*EE*_ *>* 2 + *J*_*II*_ (or equivalently 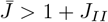). Therefore, the D domain is obtained by the unification of the two conditions

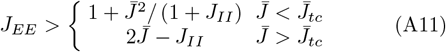

where 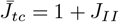.

In the case where *g* = *ge*^*±iφ*^ ∈ ℂ, that is *J*_*EE*_ *<* 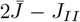, a Hopf bifurcation; i.e., *λ* = *iω* with *ω* ∈ ℝ, emerges from eq. (A9) when

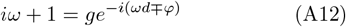

which yields two eigenvalue equations

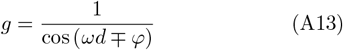

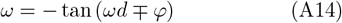

where 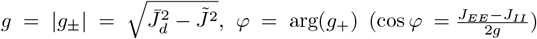 and 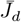 is the value of 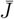 on the bifurcation line. The set of equations with − *φ* has a solution at *ω*_*d*_ ∈ [0, *π/*2*d*] (note that −*ω*_*d*_ is the solution to the set of equations with +*φ*). The Hopf bifurcation is terminated, *ω* → 0, at 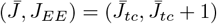.

A different Hopf bifurcation occurs for *g*_*±*_ ∈ ℝ, yielding

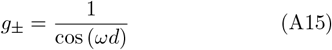

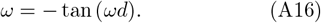

Thus, we obtain oscillations with a similar frequency as in the case of the oscillations that emerge from the OI state; compare eq. (A4) and eq. (A16). These oscillations are characterized by a fast frequency (*ω* ∈ [*π/*2*d, π/d*]). For example, in our numerical analysis *f* = *ω/*2*π* ≈142Hz for typical values *τ*_*m*_ = 2.5*d* = 5ms. Since our focus was on rhythmic activity emerging from the interplay of the E and I activities, we restricted our study to moderate values of I-to-I connectivity, *J*_*II*_ *<* 1, in which these fast oscillations do not occur. In addition, for convenience we also took *J*_*EE*_ *> J*_*II*_.

### 2. The homogeneous oscillatory solution on the bifurcation line

In the case of positive input to the inhibitory population, the two-dimensional dynamics described in eq. (2) can be reduced to a one-dimensional dynamics. This reduction is done by taking the time derivative of the equation for 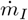 and substituting terms containing *m*_*E*_ with terms that are expressed solely by *m*_*I*_. Introducing the rescaling

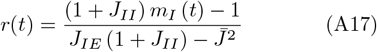

the dynamics can be rewritten in terms of the variable *r*(*t*) as

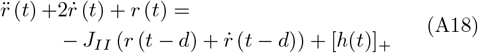

Where

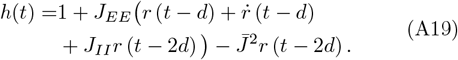

Substituting the rescaled expression for *m*_*I*_ (*t*) into its corresponding dynamical equation in eq. (2), *m*_*E*_(*t*) can be expressed in terms of *r*(*t*). This yields

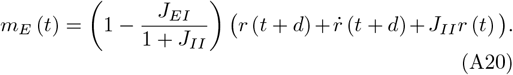

The transition from a fixed point to rhythmic activity occurs via a degenerate Hopf bifurcation. On the bifurcation line the dynamics of *r*(*t*), eq. (A18), is linear as long as *h*(*t*) is positive throughout the cycle. The degenerate solutions are given by 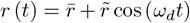, where 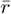 is given by the fixed point solution of eq. (A18) on the bifurcation line:

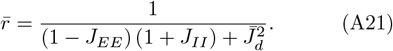

The oscillation amplitude of the degenerate solutions, 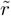, can have any value in 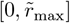, where 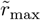 is determined by the condition that *h*(*t*) is positive to ensure linear dynamics, yielding

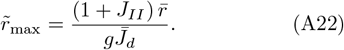

The value of 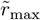 is also the oscillations amplitude in the R region immediately after the bifurcation.

The homogeneous firing rates, *m*_*I*_ and *m*_*E*_, can be obtained from *r*(*t*) using eqs. (A17) and (A20). Consequently, on the bifurcation line we can write 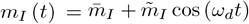 and 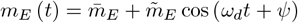, with

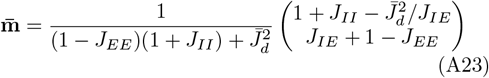

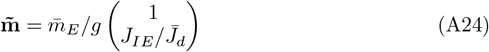

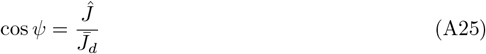

where *Ĵ* ≡ (*J*_*EE*_ + *J*_*II*_) */*2. Note that 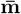 equals the fixed point **m**^*∗*^ on the bifurcation line, i.e., with 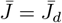.

### 3. Perturbative approach in the Fixed-Point domain

It is convenient to represent the firing rates of a network by an *N* = *N*_*E*_ + *N*_*I*_ dimensional vector 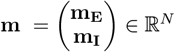 where 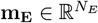 and 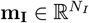 represent the activity of the excitatory and inhibitory populations, respectively. The neural dynamics, eq. (1), is now written as

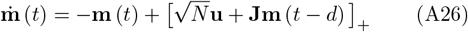

where the connectivity matrix, **J** ∈ ℝ*N ×N*

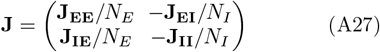

consists of the intra-connection matrices 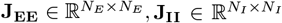 and the inter-connection matrices 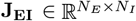 and 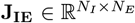. The external input to the entire population is represented by 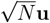 where **u** ∈ ℝ*N* is a uniform normalized vector; i.e. 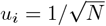.

The fixed point solution to eq. (A26) is

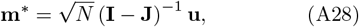

where **I** ∈ ℝ*N ×N* is the identity matrix. The connectivity matrix can be written as **J** = ⟨**J**⟩ + *δ***J** where ⟨ **J**⟩ is the homogeneous matrix ⟨*J*_*XY*_⟩ _*ij*_ = *J*_*XY*_ (*X, Y* = *E, I*), and *δ***J** is the fluctuations. Using this notation, the firing rates, **m**^*∗*^, are given by

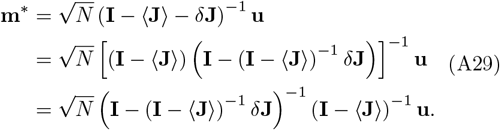

For small fluctuations, we can expand **m**^*∗*^ to a leading order in *Δ***j**

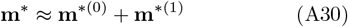

Where

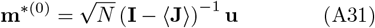

is the homogeneous fixed point solution, and

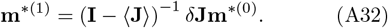

Note that *δ***Jm**^*∗*(0)^ is an eigenvector of ⟨**J**⟩ with zero eigenvalue. Consequently (**I** − ⟨**J**⟩)^*−*1^*δ***Jm**^*∗*(0)^ = **I***δ***Jm**^*∗*(0)^, yielding

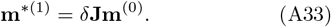

Here we only considered variability in the interconnections; i.e., *δJ*_*XX,ij*_ = 0; hence, the first order term of 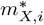 is

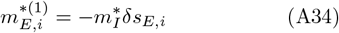

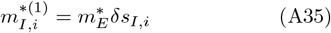

Where 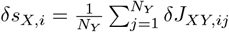, and *X* ≠*Y*. To a leading order in the fluctuations we obtain eqs. (8) and (9).

### 4. Perturbative approach in the R domain

In critical rhythmogenesis, once the system reaches the R domain it will remain near the bifurcation line. Consequently, to analyze the STDP dynamics in the R domain, we can approximate the neuronal responses by their linear dynamics on the bifurcation line. On the bifurcation line, the firing rate of neuron *n* in population *Z* is given by 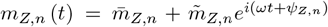, where for mathematical convenience we used complex notation for *m*_*Z,n*_(*t*) with the understanding that the firing rates are given by the real part. For small fluctuations in the synaptic weights, the firing rates can be expanded around the spatially homogeneous firing rates (i.e. for **J** = ⟨**J**⟩) to first order in *δ***J**, writing *m*_*Z,n*_ (*t*) ≈ *m*_*Z*_ (*t*) + *δm*_*Z,n*_ (*t*) where

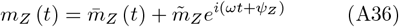

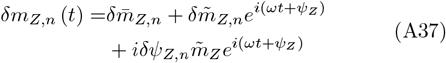

where *ψ*_*I*_ = 0, *ψ*_*E*_ = *ψ*. The fluctuations in the temporal means, 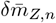, are given by eqs. (A34) and (A35).

The time dependent fluctuations, 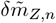 and *δψ*, are given by

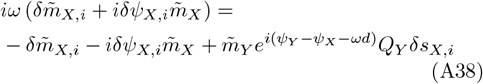

where 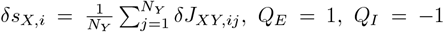, and *X* = *Y*. Solving the two equations for the real and imaginary parts of eq. (A38) we obtain

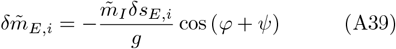

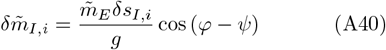

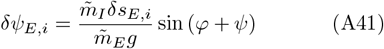

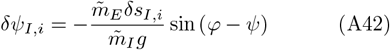

where *ψ* is given by eq. (A25). Using eqs. (A39) to (A42), to a leading order in the fluctuations *δJ*_*UV,kl*_, we obtain eqs. (10) and (11).

### 5. Critical point and conditions of critical rhythmogenesis

In order to identify the critical point on the bifurcation line 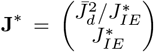, at which the drift vanishes, the vectors 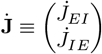 from both sides of the bifurcation line must be oriented in opposing directions. This condition can be mathematically expressed as 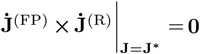, which results in

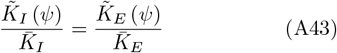

and reveals the critical point

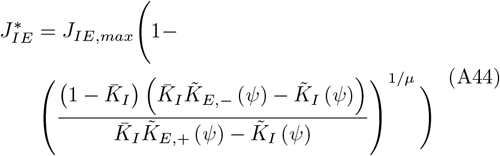

which is determined by the STDP parameters.

Two essential conditions must be met for critical rhythmogenesis to occur. First, the left branches of the *J*_*XY*_ nullclines must exhibit stability in the *J*_*XY*_ direction. Second, the excitatory nullcline must diverge from the bifurcation line at a point higher than the divergence of the inhibitory nullcline. As demonstrated in fig. 3a, when these conditions are met, the flow between the nullclines facilitates a return to the bifurcation line, thereby ensuring its stability and enabling critical rhythmogenesis (see Socolovsky and Shamir [43] for details in the case of only E-to-I and I-to-E synapses). If the excitatory nullcline diverges from the bifurcation line below the inhibitory nullcline, critical rhythmogenesis is unstable. In the limiting case where the nullclines diverge from the bifurcation line at the same point, **J**^*∗*^ coincides with the point of divergence. Therefore, the critical values of the STDP parameters that give rise to critical rhythmogenesis occur when 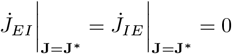. This imposes an additional condition on eq. (A43)

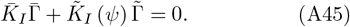

### 6. STDP dynamics of the fluctuations

The STDP dynamics for a synaptic weight *J*_*XY,ij*_, eq. (A7), is given by

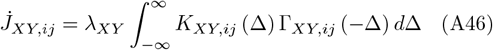

where *K*_*XY,ij*_ (Δ) = *f*_*XY*_ (*J*_*XY,ij*_) *K*_*XY*,+_ (Δ) − *αK*_*XY,−*_ (Δ) and *X* ≠ *Y*.

Considering small deviations from the homogeneous solution, *δJ*_*XY,ij*_ = *J*_*XY,ij*_ − *J*_*XY*_, we can expand eq. (A46) to the first order in these deviations. In the FP domain we obtain

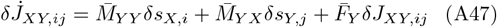

and in the R domain near the bifurcation line

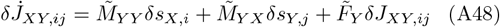

where: 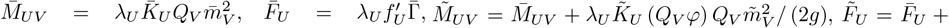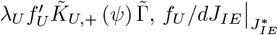, *Q*_*E*_ = 1 and *Q*_*I*_ = −1.

Eqs. (A47) and (A48) describe the linear dynamics of the fluctuations, which are characterized by four families of eigenvectors. Eigenvectors of family I represent fluctuations in the inhibitory synapses to an excitatory cell *k* (*k* ∈ 1, …, *N*_*E*_), such that ∑_*l*_ Δ*J*_*EI,kl*_ = 0. These eigenvectors are characterized by zero eigenvalues, 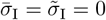, where we denoted by 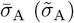 the eigenvalues of family A∈ {I, II, III, IV} in FP (R) region. The multiplicity of this family is *N*_*E*_ (*N*_*I*_ − 1).

Eigenvectors of family II represent fluctuations in the excitatory synapses to an inhibitory cell, *k* (*k* ∈ {1, … *N*_*I*_}), such that ∑_*l*_ Δ*J*_*IE,kl*_ = 0. The eigenvalues of these eigenvectors in the FP and R regions are 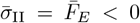 and 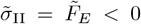, respectively. The multiplicity of this family are *N*_*I*_ (*N*_*E*_ − 1).

Eigenvectors of family III represent fluctuations in the firing of excitatory neurons such that a constant mean excitatory activity, *m*_*E*_(*t*) = ∑_*i*_ *m*_*E,i*_(*t*), is maintained. In these eigenvectors, all the inhibitory connections to each excitatory neuron *k* ∈ {1, … *M*_*E*_} are modified by the same amount, 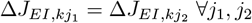, and all of the connections of excitatory neuron *k* to the inhibitory neurons obey 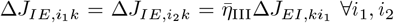 in the FP region, and 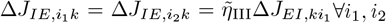 in the R region. The ratios in synaptic fluctuation are given by 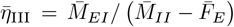 and 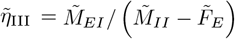. The eigenvalues of these eigenvectors in the FP and R regions are negative, 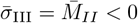 and 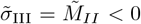. Eigenvectors in this family are determined by the deviations of the excitatory activities from the mean excitation, **m**_*E*_ − *m*_*E*_. Thus, from the condition of fixed mean activity, ∑_*i*_ *m*_*E,i*_ = *const*, the multiplicity of this family is *N*_*E*_ − 1.

Eigenvectors of family IV represent fluctuations in the firings of individual inhibitory neurons such that the mean inhibitory activity, *m*_*I*_ (*t*) = ∑_*i*_ *m*_*I,i*_(*t*), remains constant. In these eigenvectors, all the excitatory connections to each inhibitory neuron *k* are modified by the same amount, 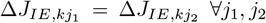, and all of the connections of excitatory neuron *k* to inhibitory neurons obey, 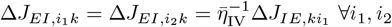 in the FP region, and 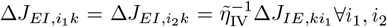 in the R region. The ratios insynaptic fluctuation are given by 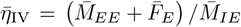 and 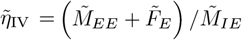. The eigenvalues of these eigenvectors in the FP and R regions are 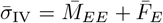 and 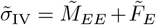, respectively. Fig. 5a depicts the eigenvalues 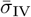 and 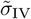 as a function of *α* for different values of *τ*_*E,−*_ (differentiated by color). As can be seen, these eigenvalues may have different signs in the FP and R regions (see section A 7 below). The multiplicity of these eigenvector s is *N*_*I*_ − 1, from the condition of fixed mean inhibition ∑_*i*_ *m*_*I,i*_ = *const*.

### 7. STDP dynamics of synaptic fluctuations in critical rhythmogenesis

Once the system reaches the bifurcation line, the order parameters *J*_*XY*_ alternate between the FP and R regions. After a transient, the order parameters will converge to alternate between asynchronous and rhythmic activities around the critical point **J**^*∗*^. The eigenvectors delineated in section A 6 result from the linearization of the STDP dynamics. Thus, in general, there are two sets of eigenvectors: one for the FP region and one for the R region.

Eigenvectors of family I (II) in the FP region are also eigenvectors of family I (II) in the R region and viceversa. In contrast, although eigenvectors of family III (IV) have a similar structure in the FP and R regions, they are not the same, since 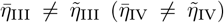. However, any eigenvector of family III (IV) from the FP region can be expressed as an eigenvector of family III (IV) from the R region plus an eigenvector of family II (I), and vice-versa. Thus, the subspaces spanned by the eigenvectors of family II and III (I and IV) in the FP and the R regions are the same. Secondly, these subspaces are invariant under the STDP dynamics that alternates between the FP and R regions.

The eigenvalues of families II and III are negative. Consequently, any fluctuation in the subspace spanned by the eigenvectors of families II and III will decay. Consider now the dynamics of a small fluctuation in the subspace spanned by the eigenvectors of family I and IV. The eigenvalue of family IV is negative in the R region and positive in the FP region. Below, we present a heuristic argument to determine whether or not fluctuations in this subspace will decay with time.

Let us denote initial conditions by **w**(0) and examine its dynamics over a full cycle of FP-R alternation. Assuming the initial state of the system is FP, it is convenient to express the initial condition by eigenvectors of the FP region: **w**(0) = *x*(0)**v**^(FP)^ + *y*(0)**u**, where **v**^(FP)^ is an eigenvector from family IV, **u** is an eigenvector of family I, and *x*(*t*), *y*(*t*) ∈ ℝ. After a duration of 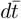 in the FP region, the fluctuation becomes 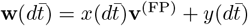, with

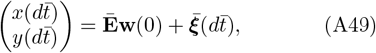

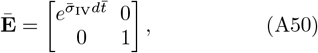

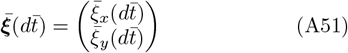

where, assuming small fluctuations, we linearized the STDP dynamics. The term 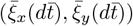 represents the projection of Gaussian white noise onto the subspace spanned by [**v**^(FP)^, **u**]. This additive white noise has zero mean, and its variance scales with the time interval 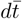; accordingly, we write 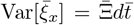 (see Appendix A 9 for details).

After transitioning to the R state at time 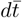 it is convenient to express the fluctuation by the eigenvectors of the R region: 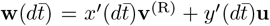, where **v**^(R)^ is the corresponding eigenvector from family IV in the R region, and *x*^*′*^, *y*^*′*^ ∈ ℝ can be obtained by a linear transformation

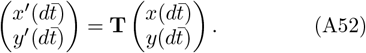

After a duration of 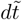in the R region the fluctuation evolves to 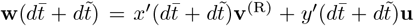, with

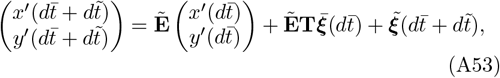

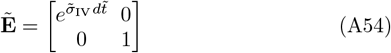

where 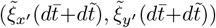 is the projection of the additive Gaussian white noise during the time interval spent in the R region, 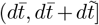, on the subspace spanned by [**v**^(R)^, **u**]. As above, the additive noise has zero mean and a variance that is proportional to the length of the time interval, 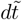; thus, we write 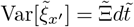.

Transitioning back to the FP region, the fluctuation 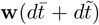 can be expressed again by **v**^(FP)^ and **u**, using the inverse transformation matrix **T**^*−*1^. Thus, after completing one full cycle

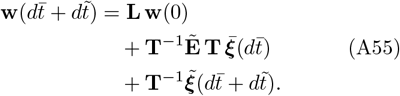

With

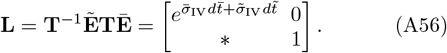

The determinant of **L**,

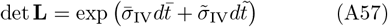

determines whether the fluctuation will grow or vanish in time. To this end, an estimation of 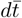 and 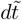 is required. The time spent in each region is determined by the dynamics of the order parameters around **J**^*∗*^. Thus, we expect 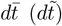 to be proportional to the inverse of the rate of exiting the FP (R) region, 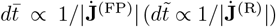.

At the critical point, 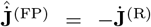 (see section III B); hence, the magnitude of the ratio of 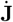 is equal to the ratio of the magnitude of each of its components, 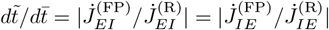. We can thus define the effective stability eigenvalue of family IV:

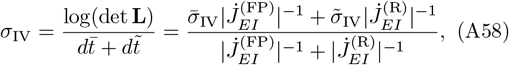

which is depicted in fig. 5b as a function of *α* for different choices of *τ*_*E,−*_ differentiated by color. Note that *σ*_IV_ is negative (stable) in the presented range.

#### Auto correlations

The fluctuations in the synaptic weights, **w**(*t*), at time *t* = *ndt* (where we denote for convenience 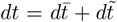) can be obtained by iteratively applying the transformation of eq. (A55) *n* times, yielding,

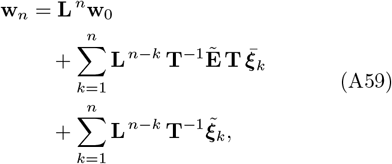

where 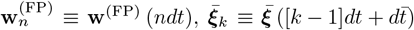and 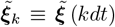. Since we are interested in the autocorrelation arising from the additive noise, it is convenient to subtract the transient term due to initial conditions and focus on 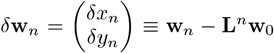. From eq. (A59) one obtains

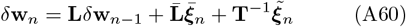

where 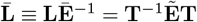.

#### Computation of the matrix

**T**. Because the eigenvectors from family I are independent of the state of the system (FP or R), **T** can be expressed as

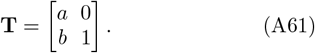

Similarly

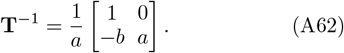

To determine *a* and *b*, we consider a simple illustrative case. Specifically, we take *N*_*E*_ = *N*_*I*_ = 3 and examine a family IV eigenvector **v**^(*β*)^ (*β* = {FP, R}) with: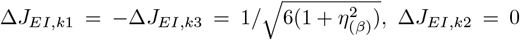 and Δ*J*_*IE,ki*_ = *η*_(*β*)_Δ*J*_*EI,ik*_ where 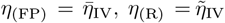 and *i, k* ∈ {1, 2, 3} (see Appendix A 6 for details on family IV eigenvectors). Thus, **v**^(FP)^ can be expressed as a linear combination of **v**^(R)^ and a family I vector, **u**, that satisfies 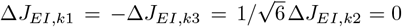and Δ*J*_*IE,ij*_ = 0 for *i, j, k* = 1, 2, 3. finally, from **v**^(FP)^ = *a***v**^(R)^ + *b***u** one obtains: 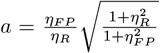and 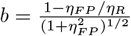.

Using the fact that the upper right elements of **T** and **T**^*−*1^ are zero, eq. (A60) yields

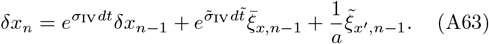

To obtain the autocorrelation, *R*_*x*_(*ndt*, [*n* + *m*]*dt*) = ⟨*δx*_*n*_*δx*_*n*+*m*_⟩, we multiply eq. (A63) at times *ndt* and (*n* + *m*)*dt* and average over the white noise statistics, yielding

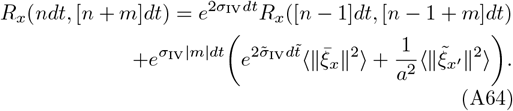

For long times (and *σ*_IV_ *<* 0) the autocorrelation converges, 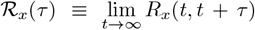, one obtains from eq. (A64) (assuming 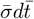 and and 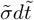 small):

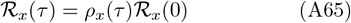

where the variance, ℛ_*x*_(0), is given by

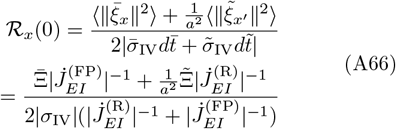

and the correlation coefficient, *ρ*_*x*_(*τ*), is given by

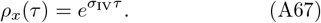

Similarly, by following the steps outlined in eqs. (A51) and (A67), the long-time autocorrelation of the synaptic weight fluctuations in the R region, is given by

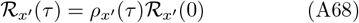

where the variance, 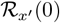, is given by

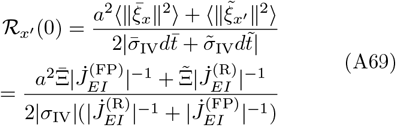

and *ρ*_*x*_*′* (*τ*) = *ρ*_*x*_(*τ*).

We can therefore define the effective autocorrelation of the fluctuations projected onto an eigenvector of family IV as

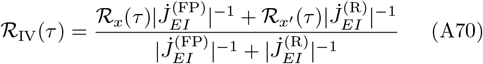

where the effective variance is given by

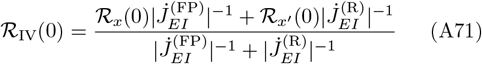

and the effective correlation coefficient is

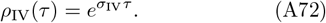

The effective variance, ℛ_IV_(0) and effective correlation coefficient *ρ*_IV_(*τ*) are depicted in dashed black lines in fig. 6a and in fig. 6b, respectively.

### 8. STDP dynamics of intra-connection order parameters

Assuming the intra-connections are plastic, the induced STDP dynamics of their order parameters, *J*_*EE*_ and *J*_*II*_, can be derived similar to eq. (A8), yielding

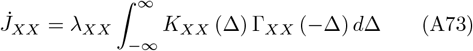

where *K*_*XX*_ (Δ) = *f*_*XX*_ (*J*_*XX*_) *K*_*XX*,+_(Δ) −*αK*_*XX,−*_(Δ) and 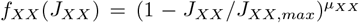. Assuming that the STDP dynamics of the inter-connections, *J*_*EI*_ and *J*_*IE*_, drive the system toward a state of critical rhythmogenesis the homogeneous firing rates, *m*_*X*_ (*t*), are given by eq. (A4) in the R region near the bifurcation line. Therefore, the autocorrelation in the state of critical rhythmogenesis takes the form

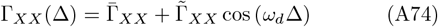

where 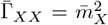 (in general in the FP region 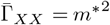). In the FP region, 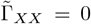, whereas in the R region, 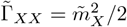. Accordingly, the STDP dynamics becomes

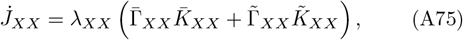

Where

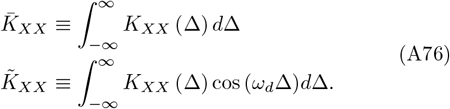

In our choice of normalization, 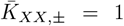, which yields 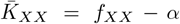. For the exponential temporally asymmetric Hebbian and anti-Hebbian learning rules (eq. (A5)), we obtain

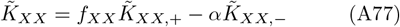

with 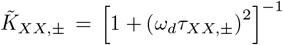. As in the case of the inter-connections, we assume *τ*_*XX*,+_ *< τ*_*XX,−*_ and *α < f*_*XX*_ *<* 1, see Appendix A 11. In the case of an additive learning rule (*µ*_*XX*_ = 0), 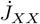 is strictly positive, driving the system toward a pathological state. To prevent this, a homeostatic plasticity mechanism must be introduced; namely, a non-additive learning rule with *µ*_*XX*_ *>* 0. Figure 8 illustrates the STDP dynamics in the plane of 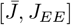, depicted by the blue dots. Starting from initial conditions in the FP region (red ‘+’) the system is drawn towards the R region where it remains near the bifurcation line alternating repeatedly between fixed point and rhythmic activity. The oscillation frequency in the rhythmic region is shown in the inset.

**FIG. 8:**
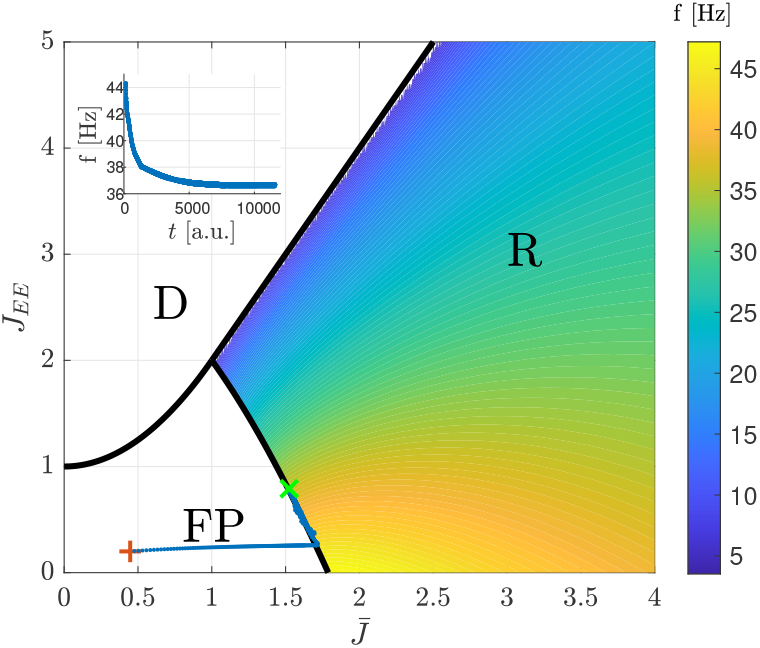
Critical rhythmogenesis with plastic *J*_*EE*_, *J*_*IE*_, and *J*_*EI*_ (See Appendix A 9 for details). The STDP dynamics of the order parameters, eqs. (18) and (A73) are shown by the blue dots in the plane of 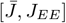 (compare with the phase diagram in fig. 2b). The trajectory starts at the red ‘+’ and ends at the green ‘*×*’. The inset demonstrates the frequency of the system when it exhibits rhythmic activity (in the R region), as a function of time. The initial conditions we chose are *J*_*EI*_ = 0.2, *J*_*IE*_ = 1, *J*_*EE*_ = 0.2 and *J*_*II*_ = 0. Here we used *α* = 0.98, *µ*_*IE*_ = 0.01, *µ*_*EE*_ = 0.02, *J*_*IE,max*_ = 20, *J*_*EE,max*_ = 1, *τ*_*E*,+_ = *τ*_*I*,+_ = 10 ms, *τ*_*E,−*_ = 25 ms, *τ*_*I,−*_ = 15 ms, and *τ*_*m*_ = 2.5*d* = 5 ms. Details on the learning rates *λ*_*XX*_ are given in Appendix A 9.

Note that the rhythm itself depends on the value of the intra-connection (see color map in fig. 8). Thus, unlike the STDP dynamics of the inter-connections, here the resultant rhythm will depend on parameters that characterize the STDP rule.

### 9. Specifics of the numerical simulations and figures

The numerical simulations in this study were performed using MATLAB, and are available for download at [78]. STDP dynamics, eqs. (17), (18) and (A73), were solved using Euler’s method for differential equations. Unless otherwise stated, the cross-correlations of the neuronal responses, Γ_*XY,ij*_(Δ) (Γ_*XY*_ (Δ)), were computed exploiting the separation of timescales in the following manner. In the FP region the correlations were computed from the exact solution of the fixed-point of the neuronal responses, given fixed synaptic weights. In the R region the correlations were computed by simulating the neuronal dynamics, eqs. (1) and (2), with fixed synaptic weights over a time interval of Δ*t* = 200*τ*_*m*_ with a time step of *dt* = 0.01*τ*_*m*_. The final period of *m*_*X,i*_(*t*) and *m*_*Y,j*_(*t*) (*m*_*X*_ (*t*) and *m*_*Y*_ (*t*)) was used to compute the cross-correlation leveraging the periodicity of Γ_*XY,ij*_(Δ) (Γ_*XY*_ (Δ)). The boundaries of the integral in the right hand side of eq. (A7) (eq. (A8)) were replaced with ±Δ*t* = ±200*τ*_*m*_ and a time bin of *dt* = 0.01*τ*_*m*_ was used for the numerical integration.

In fig. 3c (and red trace in fig. 3b) 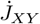 was estimated from the continuous time dynamics of the neuronal responses:

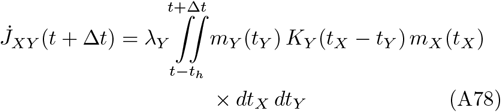

where a history trace up to *t*_*h*_ = 50*τ*_*m*_ was used and *K*_*Y*_ (*t*_*X*_ − *t*_*Y*_) = *f*_*Y*_ (*J*_*XY*_)*K*_*Y*,+_(*t*_*X*_ − *t*_*Y*_) −*αK*_*Y,−*_(*t*_*X*_ − *t*_*Y*_).

In fig. 7, three inhibitory neurons were removed at time *t*_0_ = 3000 [a.u.] from a network initially consisting of *N*_*E*_ = 40 excitatory and *N*_*I*,initial_ = 10 inhibitory neurons. The simulation was continued until time *t* = 8000 [a.u.]. Time is expressed in units of the integration time step, Δ*t*, used in the numerical simulation of the STDP dynamics.

In fig. 7, the mean frequencies during critical rhythmogenesis (before and after *t*_0_) were obtained by averaging the network’s observed frequencies across all instances in which the system was in the R region. The corresponding uncertainties were quantified as the standard deviation of these frequencies.

In fig. 8, two sets of learning rates were employed. When the STDP dynamics slowed down, specifically when 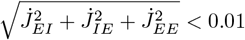, the learning rates were set to 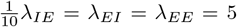 to accelerate the simulations. Otherwise, the learning rates remained at their baseline values, 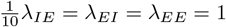.

In figs. 6 and 7 Gaussian white noise was incorporated into the simulated dynamics of the synaptic plasticity:

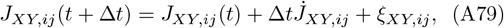

where 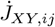 is given by eq. (A7), and *ξ*_*XY,ij*_ represents independent Gaussian white noise. Specifically, the noise terms were drawn from *ξ*_*EI,ij*_ ∼ *N* (0, 10^*−*4^) and *ξ*_*IE,ij*_ ∼ *N* (0, 10^*−*2^).

### 10. Diluted connectivity

We simulated the STDP dynamics in a diluted connectivity network, fig. 9. Initially, synapses were removed through an i.i.d. process with survival probability *p*. Under this fixed diluted architecture, the STDP dynamics of the remaining synapses of the inter-connections was simulated from the continuous time dynamics of the neuronal responses:

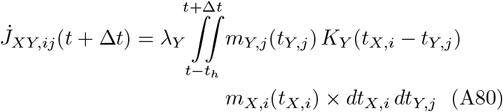

where a history trace up to *t*_*h*_ = 50*τ*_*m*_ was used and *K*_*Y*_ (*t*_*X,i*_ − *t*_*Y,j*_) = *f*_*Y*_ (*J*_*XY,ij*_)*K*_*Y*,+_(*t*_*X,i*_ − *t*_*Y,j*_) − *αK*_*Y,−*_(*t*_*X*.*i*_ − *t*_*Y,j*_). The STDP dynamics drive the order parameters *J*_*XY*_, after a brief transient, toward critical rhythmogenesis on the bifurcation line (trajectory depicted with blue dots).

**FIG. 9:**
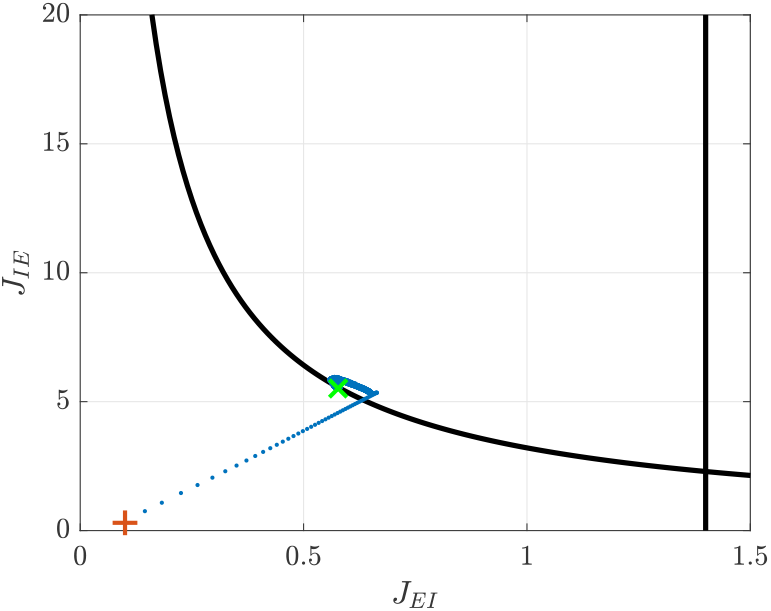
Diluted connectivity. STDP dynamics of the inter-connections was simulated in a diluted network of *N*_*E*_ = 60 excitatory and *N*_*I*_ = 15 inhibitory neurons and dilution parameter *p* = 0.7 was simulated. Note that all synapses were subject to the dilution process, see Appendix A 10. The initial conditions for the synaptic weights of the remaining inter-synapses were drawn independently from a Gaussian distribution, *J*_*XY,ij*_ *∼ N* (*J*_*XY*,0_, [*J*_*XY*,0_*/*10]^2^), with *J*_*EI*,0_ = 0.1 and *J*_*IE*,0_ = 0.3. The synaptic weights of the remaining intra connections were set to *J*_*EE,ij*_ = 0.6 and *J*_*II,ij*_ = 0.4. The trajectory of the order parameters is shown in the [*J*_*EI*_, *J*_*IE*_] plane with the blue dots. The initial condition is marked by the red ‘+’ and the green ‘*×*’ marks the location at the end of the simulation. Here we used *α* = 0.98, *µ* = 0.01, *λ*_*E*_ = 10*λ*_*I*_ = 0.8 *J*_*IE,max*_ = 20, *τ*_*E*,+_ = *τ*_*I*,+_ = 10 ms, *τ*_*E,−*_ = 25 ms, *τ*_*I,−*_ = 15 ms, and *τ*_*m*_ = 2.5*d* = 5 ms002E

In order to maintain the same phase diagram, the normalization by the population sizes *N*_*E*_ and *N*_*I*_ in neuronal response dynamics, eq. (1), was replaced with normalization by the mean number of synaptic connections, namely *pN*_*E*_ and *pN*_*I*_.

### 11. Choice of Parameters

The key parameters used in this study were chosen based on a combination of theoretical and experimental considerations, as detailed below:

- Due to the strong positive feedback inherent in STDP dynamics, the relative strength of depression, *α*, is typically set close to 1 to balance potentiation and depression (see, e.g., Ref. [36]). In our simulations, we used *α* values in the range [0.9, 1].
- Experimental studies show temporal kernels of STDP with time constants that are on the order of several tens of milliseconds [33, 34, 39, 40, 49, 79, 80]. As in [34, 39, 49], we focused on temporally asymmetric STDP rules with *τ*_+_ *< τ*_*−*_.
- The ratio of excitatory to inhibitory learning rates, *λ*_*E*_*/λ*_*I*_, was chosen to reflect the typical dynamic ranges of the synaptic weights; for instance, as illustrated in Fig. 2a, *J*_*IE*_ spans roughly 10, while *J*_*EI*_ spans about 1.
- Noise in the STDP dynamics may arise from several sources. One contribution stems from the intrinsic variability of neuronal responses, which drives the STDP dynamics. However, in the limit of slow learning, *λ* → 0, this component vanishes under the mean-field Fokker–Planck approximation [36]. A second potential source is activity-independent synaptic motility [4, 5, 7, 8, 81–83]. For example, Shomar et al. reported fluctuations of approximately 30% over a 24-hour period [8]. Nevertheless, a reliable estimate of the magnitude of synaptic-weight fluctuations on the timescales relevant to this study—seconds to minutes—remains to be desired.

In our simulations, noise was introduced primarily to probe the stability of the STDP dynamics rather than to model a specific biophysical noise source. Accordingly, we did not examine regimes of strong noise. Yet, the standard deviation of the white-noise term *ξ*_*XY,ij*_ in eq. (A79) was scaled so that the excitatory-to-inhibitory ratio was set to 10:1, consistent with the typical dynamic range of excitatory and inhibitory synaptic weights in the simulations.

## ACKNOWLEDGMENTS

This research was supported by the Israel Science Foundation (ISF) grant numbers 300/16 and 624/22.

